# Synchronized Seasonal Shifts in the Climate, Diet, and Gut Microbiome of a Rainforest Lemur (*Eulemur rufifrons*)

**DOI:** 10.64898/2026.11.18.767713

**Authors:** Jaesylin N. Stephens, David W. Weisrock, Jean Patrique Raheimanjato, Patricia C. Wright, Mariah E. Donohue

## Abstract

Gut microbiome (GM) flexibility extends the adaptive potential of host species in the face of rapid ecological change. Madagascar’s endemic primates offer a powerful natural system to study GM plasticity as the island experiences substantial environmental change throughout the year. This is especially true in the eastern rainforests, which experience marked variation in temperature, rainfall, and fruit availability between the wet and dry seasons. To elucidate interactions between climate, diet, and the GM, we conducted a 10-month study of the ecologically flexible red-fronted brown lemur (*Eulemur rufifrons*) in Ranomafana National Park. We detected significant shifts in gut microbial communities correlated with monthly rainfall, temperature, and dietary diversity. Trade-offs between abundances of carbohydrate- and fiber-metabolizing bacteria reflected shifts between frugivory and folivory, suggesting that the GM buffers energetic and/or nutritional imbalances when resources are scarce. Finally, we observed synchronized shifts in climate and the GM despite deviation from normal seasonal patterns, demonstrating the GM’s rapid response to environmental cues. Our results suggest that the GM contributes to phenotypic plasticity across seasons, enabling endangered primates to better cope with environmental stressors like climate change.

## 1 Introduction

Primates are highly attuned to seasonal climate patterns that govern the timing and accessibility of essential resources, foraging opportunities, and suitable habitats (Nagahara et al., 2025; Ozgul et al., 2023; Xue et al., 2023). Climate change, deforestation, and other anthropogenic disturbances are increasingly disrupting these patterns, making ecological flexibility critical for species persistence in the Anthropocene. This is particularly important given that the rate of Primate niche evolution cannot keep pace with climate change projections (Meyer and Pie, 2022).

The gut microbiome (GM) may play a key role in shaping Primate ecological flexibility (Henry et al., 2021). Highly responsive to changes in the host’s external environment, the GM contributes to physiological and metabolic adjustments that enable hosts to cope with fluctuating environmental conditions (Khakisahneh et al., 2020; Moeller and Sanders, 2020). This dynamic microbial community can undergo rapid compositional and metabolic changes within days or even hours of exposure to new stimuli – most notably, diet (David et al., 2014; Letourneau et al., 2022). This plasticity may offer new combinations of microbial genes that expand host metabolic capabilities, helping meet nutritional and energetic demands (Marsh et al., 2024; Moeller and Sanders, 2020). Generalist host species may especially benefit from GM plasticity as they must regularly modify foraging strategies (Marsh et al., 2024). Since GM services have been essential in the evolution of herbivory, host diet and GM composition are inextricably linked.

Species adapted to seasonal habitats exhibit predictable GM shifts throughout the year. Dry-to-rainy season GM cycles have been observed in many wild primates, including great apes (Hicks et al., 2018), chlorocebus monkeys (Trosvik et al., 2018), capuchin monkeys (Orkin et al., 2019), Ethiopian geladas (Baniel et al., 2021), Tibetan macaques (B. Sun et al., 2016; Xia et al., 2021), and several lemur species (de Winter et al., 2020; Springer et al., 2017). Variability in rainfall and, to a lesser extent, temperature may indirectly shape GM composition by altering diet, water availability, and even sociality (Beeby et al., 2025). In geladas, cumulative rainfall was a major predictor of over 60% of bacterial families in the gut (Baniel et al., 2021). Microbial loads in watering holes may increase during drier periods (Murillo, Schneider, Fichtel, and Daniel, 2022), increasing horizontal transmission of microbes that colonize the gut. However, seasonal diet is likely the strongest driver of seasonal GM cycling. A landmark human study linked GM cycling in Hadza hunter-gatherers with seasonal shifts between fiber-rich tubers and foraged berries, while industrialized populations with more uniform diets had lower abundances of seasonally fluctuating gut microbiota (Smits et al., 2017). Understanding how GMs enable hosts to cope with seasonal variation in real time is important for understanding host adaptation in unstable environments. However, more studies are needed to capture the range, speed, and function of seasonal shifts in the GM, especially during times of food scarcity.

Located in southern tropical latitudes, Madagascar’s endemic lemurs are subject to a temporally variable climate: a warm, rainy season followed by a cool, dry season. The island harbors further climatic variation geographically as its eastern coast experiences hotter temperatures, more rainfall, and more frequent cyclones compared to its western coast (Gouveia et al., 2024; Mandl et al., 2018). *Eulemur rufifrons* is a widespread species of brown lemur that spans several climate zones from coast to coast (Blair et al., 2013, Figure 1). While western populations occupy a dry deciduous forest with moderate changes in temperature and rainfall, eastern rainforest populations are faced with extreme heat and heavy rain (Donohue et al., 2022). The feeding ecology of rainforest-dwelling lemurs is determined by ripe fruit availability dependent on rainfall (Beeby and Baden, 2021; Wright, 1999). This poses a particular challenge for frugivorous eastern *E. rufifrons*, which specialize on abundant fruits during the wet season, then resort to hard-to-digest fallback foods (i.e., leaves, seeds, and flowers) when fruit is scarce during the dry season (Murillo, Schneider, Fichtel, and Daniel, 2022; Sato et al., 2016). Thus, the high degree of climatic and dietary variability *E. rufifrons* experiences annually makes it a good study system for examining GM plasticity in response to the host environment.

**Figure 1.**
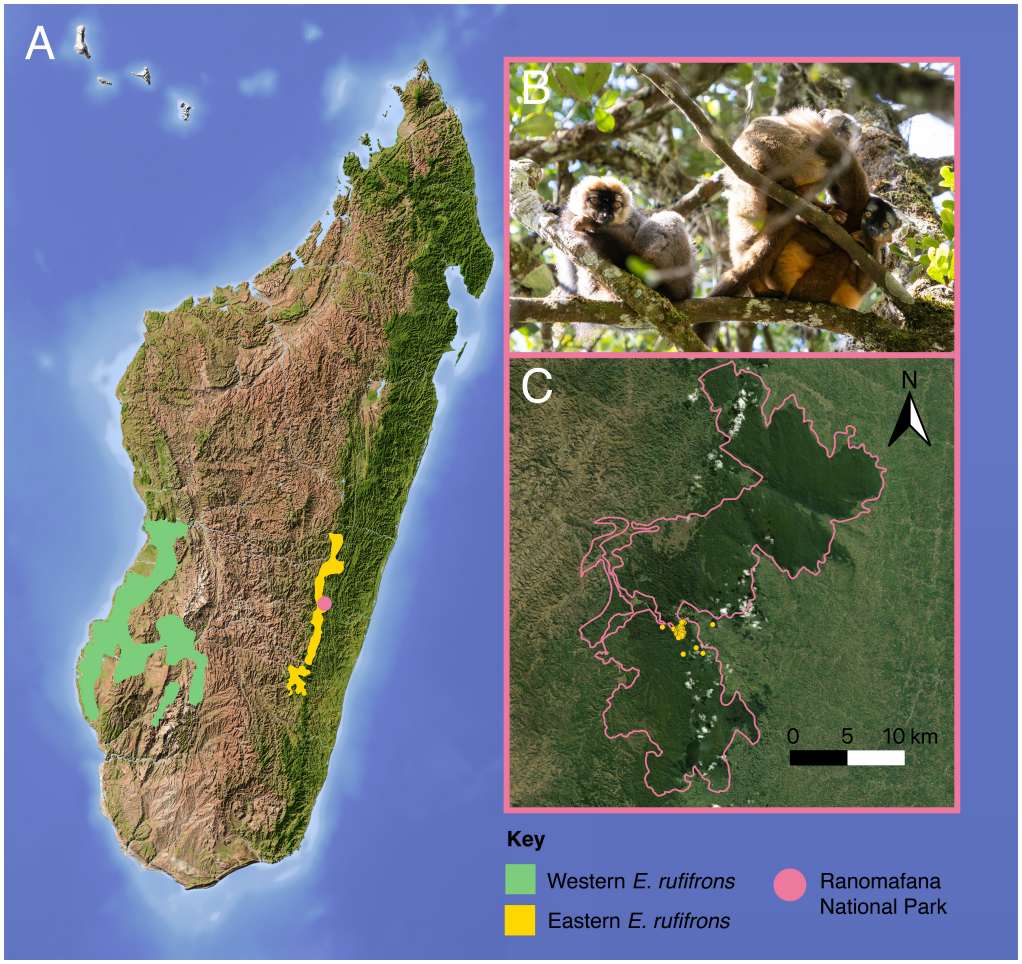
Study population and sampling location of red-fronted brown lemurs (*Eulemur rufifrons*). **A)** The geographical range of eastern (rainforest) versus western (dry forest) populations of *E. rufifrons* in Madagascar, based on IUCN range data (Johnson et al., 2020). **B)** Eastern *E. rufifrons* in Ranomafana National Park (RNP): adult male (left), juvenile (upper right), and adult female (lower right); photograph taken July 2025. **C)** Sampling sites (yellow points) within the RNP boundary (pink outline) mapped from the UNEP-WCMC/IUCN World Database on Protected and Conserved Areas (2026).

We investigated fine-scale microbiome fluctuations across ten months in *E. rufifrons* of Ranomafana National Park (RNP), a 41,000 hectare protected tract of submontane rainforest in southeastern Madagascar (Wright et al., 2012). From November to April, RNP experiences a hot, wet season with approximately 3,000 millimeters (mm) of rain and high fruit availability. May to October is traditionally considered the cool, dry season with less rain and fruit (Beeby and Baden, 2021; Wright et al., 2012). Previously, GM seasonality has been detected in dry forest lemurs, including western *E. rufifrons* (de Winter et al., 2020; Murillo, Schneider, Heistermann, et al., 2022; Springer et al., 2017), but much less is known about these dynamics in rainforest-dwelling lemurs. Given that they experience more pronounced seasonal changes than dry forests, we expect more dramatic microbiome responses in rainforest habitats. Using 16S rRNA metabarcoding data, we tested a general model of seasonal microbiome plasticity in which we predicted (1) differences in climate, diet, and gut microbiota between the dry and wet seasons, (2) a relationship between dietary and GM diversity, and (3) variation in microbial metabolic pathways that reflects dietary macronutrients. This work contributes to a growing body of research highlighting GM plasticity as a key source of adaptive potential that enhances species survival in the Anthropocene.

## 2 Methods

### 2.1 Ethics Statement

All field methods and sample collection were conducted noninvasively according to methods approved in a University of Kentucky Institutional Animal Care and Use Committee (IACUC) protocol (IACUC #2018-2919 and #2021-3814), with field methods approved by Madagascar National Parks permit #132/22/MEDD/SG/DGGE/DAPRNE/SCBE.Re, and in adherence with the American Society of Primatologists (ASP) Principles for the Ethical Treatment of Non-Human Primates.

### 2.2 Field Methods

We conducted full-day focal follows of *E. rufifrons* in RNP during two field seasons (January – February and July – October, 2020; Table S1). Focal animal behaviors were recorded instantaneously every three minutes on days that fecal samples were collected, culminating in 165 total hours of behavior data. Activity categories consisted of feeding, moving within trees, resting, traveling between trees, and social behaviors (grooming, playing, fighting, vocalizing, and tail wagging). We estimate that our data include at least five social groups, though the identities of groups and individuals were not always certain as only one group was collared. We also recorded foods consumed during behavioral observations. Plant species were initially identified to their Malagasy vernacular name, and later matched with scientific name whenever possible. All food resources were categorized as fruit, leaves, flowers, seeds, or soil.

We opportunistically collected 61 fecal samples (*n*_*wet*_ = 33, *n*_*dry*_ = 28) from one field site (Talatakely). To prevent contamination of soil microbes, the center of feces was collected with sterile tweezers within one minute of deposition. Samples were aseptically transferred to tubes of 96% ethanol and stored at room temperature until laboratory processing.

A map of the study area in RNP was produced using QGIS (v. 4.2; QGIS Development Team, 2026). Coordinates were recorded during field sampling and referenced to the World Geodetic System 1984 (WGS 84; EPSG:4326). The boundary of RNP (protected-area identifier 20287; updated 2026) was obtained from the World Database on Protected and Conserved Areas (WDPCA), maintained by the United Nations Environment Programme World Conservation Monitoring Centre (UNEP-WCMC) and the International Union for Conservation of Nature (IUCN).

### 2.3 Climate Analysis

Climate data included daily cumulative rainfall and air temperature measurements taken daily at 6:00 AM at Centre ValBio research station in RNP between January 1, 2020 and October 31, 2020. These data were provided by The Institute for the Conservation of Tropical Environments (ICTE) at Stony Brook University. Daily measurements were taken using a rain gauge and a thermometer with minimum temperature (*T*_*min*_)and maximum temperature (*T*_*max*_) for the last measurement period. To maintain consistency with GM data, we only analyzed climate data for months that fecal samples were collected. We defined seasonal cutoffs according to conventional monthly classifications with the exception of October, which this study includes as a wet season month based on 2020 climate patterns. We created two separate datasets for each climate variable to make comparisons between the wet and dry seasons and between individual months. All analyses and plots were generated in R.

For rainfall data, we calculated descriptive statistics (mean, standard deviation, median, and interquartile range) and cumulative rainfall for each season and month. Potential outliers were retained in the analyses. We performed nonparametric tests to compare rainfall between each month and between the wet and dry seasons due to non-normally distributed data. We compared seasons using a two-sided Wilcoxon rank-sum test (Mann-Whitney U test). We compared daily cumulative rainfall measurements between months with Kruskal-Wallis rank-sum tests. Significant overall month effects were followed by pairwise comparisons using Dunn’s post-hoc tests. *p*-values were adjusted using the Holm procedure to control for the family-wise error rate.

For temperature data, we calculated mean daily temperature by averaging *Tmin* and *Tmax*. We calculated descriptive statistics by season and by month. Seasonal differences were tested with a Welch’s unpaired t-test on seasonal means, while monthly differences were tested with Kruskal-Wallis tests on monthly medians. A significant omnibus test was followed by pairwise month comparisons using Dunn’s tests. *p*-values from pairwise tests were adjusted using the Holm procedure.

### 2.4 Diet and Behavior Analysis

Separate baseline-category multinomial mixed-effects logistic regression models were fitted to test variation in diet composition among months and between seasons using the `mblogit()` function in the mclogit R package. The models accounted for non-independence among observations collected during the same focal-follow day by including observation data as a random effect. Fruit was the reference response category.

Overall month and season effects were evaluated using joint Wald chi-square tests across the multinomial response. Pairwise tests were performed for monthly comparisons, with *p*-values adjusted using the Holm method for multiple comparisons. For descriptive summaries, category counts were pooled within each month and season, expressed as percentages of all observations in that period, and plotted. Daily food proportions were calculated and plotted to display variation among observation days. The same multinomial model and descriptive statistics approaches were used to test monthly and seasonal differences in activity categories, but with resting as the reference response category. We note that October was excluded from these analyses due to lack of behavioral data.

### 2.5 DNA Sequence Generation

16S rRNA (v4 region) sequence data were generated from 61 fecal samples. Samples were randomized during the extraction and PCR steps to control for batch effects. DNA was extracted from ∼0.2 g of each fecal sample using the Qiagen QIAamp® PowerFecal® Kit per the manufacturer’s protocol. PCR amplification was performed using unique dual-indexed primers (Kozich et al., 2013), in a reaction volume containing 22.5 µL of Accuprime Pfx Supermix™, 4 µL of DNA, and 1 µL each of a forward and reverse primer (10 µM concentration). Amplification occurred in a thermal cycler with the following touchdown protocol: 2 min. at 95°C (initial denaturation), 19 cycles of 20 sec. at 95°C, 15 sec. at 60°C, 5 min. at 72°C, 19 cycles of 20 sec. at 95°C, 15 sec. at 55°C, 5 min. at 72°C, and 10 min. at 72°C. PCR products were confirmed with agarose gel electrophoresis. PCR products were quantified with a Qubit fluorometer. DNA concentration was normalized to 1 ng/µL, then pooled. Paired-end sequencing (2x250bp) was performed at the UK HealthCare Genomics Core on an Illumina MiSeq flow cell.

### 2.6 Bioinformatics

Raw Illumina sequence processing and bioinformatic analyses were conducted using QIIME 2 (v. 2022.2; Bolyen et al., 2019). Sequencing reads were filtered and denoised with the DADA2 pipeline (Callahan et al., 2016) and truncated to a maximum of 230 bp. Rooted microbial phylogenetic trees were generated for downstream core diversity analyses using the `align-to-tree-mafft-fasttree` pipeline. Data were rarefied to 9,700 reads, the dataset’s smallest library size above 1,000 reads, to standardize sequencing depth across samples and maximize sample retention. Microbial taxonomy was assigned using 99% clustering to the SILVA reference database (Quast et al., 2013).

Following quality filtering and taxonomic assignment, an amplicon sequence variant (ASV) abundance table was generated. For each sample, the relative abundance of each taxon was calculated by dividing its read count by the total number of reads at that taxonomic level, expressed as percentages.

First, we asked whether GM composition could accurately predict sampling season using QIIME 2’s q2-sample-classifier. This analysis uses a random forest machine learning algorithm to predict metadata assignment of test samples by learning the microbial composition of training samples with provided metadata (Bokulich et al., 2018). The top 50 most informative ASVs were visualized with a heatmap.

Next, we evaluated sources of variation in microbiome alpha diversity. We performed Kruskal-Wallis tests to determine how sampling season, month, host sex, and host individual impacted three measures of alpha diversity: Faith’s Phylogenetic Diversity (PD), Shannon Diversity Index, and Observed Operational Taxonomic Units (OTUs). We used Linear Mixed Models (LMMs) with the R package lme4 (v. 1.1-37) (Bates et al., 2015) to assess the relationship between monthly climate variables (cumulative rainfall (mm), minimum temperature, and maximum temperature (°C)) and alpha diversity. A separate LMM and plot of the fixed effect model was generated for each climate variable and alpha diversity metric. We included lemur ID as a random effect. We visually checked each model’s residuals for deviation from a normal distribution and heteroscedasticity.

We quantified variation in GM community composition using four measures of beta diversity: Bray–Curtis dissimilarity, Jaccard distance, weighted UniFrac, and unweighted UniFrac. Bray–Curtis and Jaccard measure similarity and dissimilarity using microbial taxonomy, whereas UniFrac incorporates phylogenetic relationships among microbial taxa and branch lengths when calculating between-community distances. Bray–Curtis and weighted UniFrac are abundance-weighted measures that factor in relative OTU abundances, whereas Jaccard and unweighted UniFrac use OTU presence–absence measures. We assessed the effects of season, month, the season × month interaction, sex, and individual on beta diversity using permutational multivariate analysis of variance (PERMANOVA) with 999 permutations and implemented with the `adonis` method (p-formula = `Season + Month + Season/Month + Sex + Individual`). Beta-diversity patterns were visualized in ordination space using principal coordinates analysis (PCoA), with ordinations generated in R using the phyloseq package (v1.50.0).

For more precise estimates of the relationship between climate and GM beta diversity, we performed principal component analysis (PCA) using the R packages compositions (v. 2.0-8) and vegan (v. 2.6-10). We calculated Aitchison distances by centered log-ratio (CLR) transforming and calculating Euclidean distance for our rarefied sample ASV counts. We generated PCA scatterplots and colored the points by rainfall or temperature. We used vegan’s `envfit()` function to project climate variables onto PCA ordination with 999 permutations. Vectors for significant variables were overlaid on the PCA plot.

To infer metabolic functional capabilities of the GM, we used PICRUSt2 to predict metagenome-based marker genes (i.e., 16S data) (Douglas et al., 2020). We normalized MetaCyc pathway counts and removed pathways with a mean relative abundance less than 0.1% across samples. Using the R package tidyverse (v. 2.0.0), we performed Wilcoxon rank-sum tests and adjusted p-values using the Benjamini-Hochberg False Discovery Rate (FDR) method to compare pathway relative abundances between seasons. To visualize seasonal differences, we calculated the group-wise mean difference (Dry - Wet) and generated diverging bar plots with a significance threshold of *p* < 0.0001, as well as *p* < 1 ×10^−9^ for optimal visualization.

All data visualizations were created using the package ggplot2 (v. 3.5.2) in RStudio (v. 2024.09.0).

## 3 Results

### 3.1 Climate

There were significant differences in rainfall and temperature patterns between the wet and dry seasons (Figure 2). A total of 183 daily rainfall and temperature observations across the six sampling months were analyzed (*n*_*wet*_ = 91, *n*_*dry*_ = 92; Table S2). Rainfall was significantly higher during the wet versus dry season, with 1,434.4 mm vs. 434.8 mm of cumulative rainfall, respectively. Mean daily rainfall was 15.8 ±20.8 mm during the wet and 4.73 ±9.9 mm during the dry season. Median daily rainfall was 7.2 mm (IQR: 0.0–25.8) during the wet and 0.0 mm (IQR: 0.0–4.9) during the dry season (Wilcoxon *W* = 5904, *p* = 6.01; Table 1 and Figure 2A). The overall month effect of daily rainfall was significant (Kruskal-Wallis 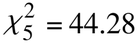, *p* = 2.04 ×10^−8^). January had the highest cumulative rainfall among sampled months (672.0 mm), whereas August had the lowest (43.2 mm). January also had the highest median daily rainfall at 18.4 mm (IQR: 2.6–31.9), while the median for August and September was 0.0 mm. Six pairwise tests remained statistically significant after Holm adjustment (Table S3 and Figure S1).

**Table 1.**
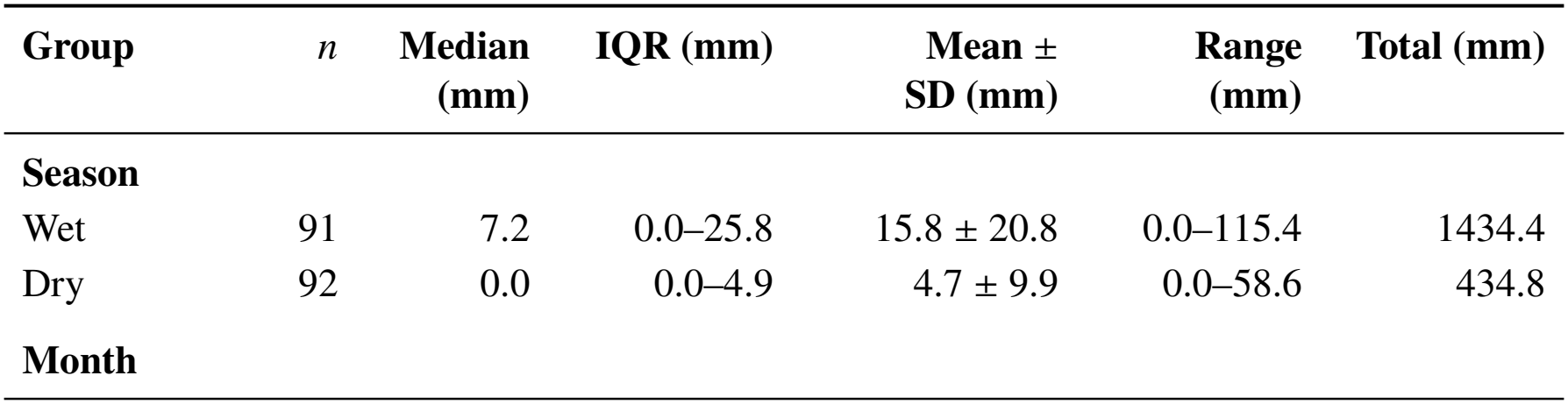
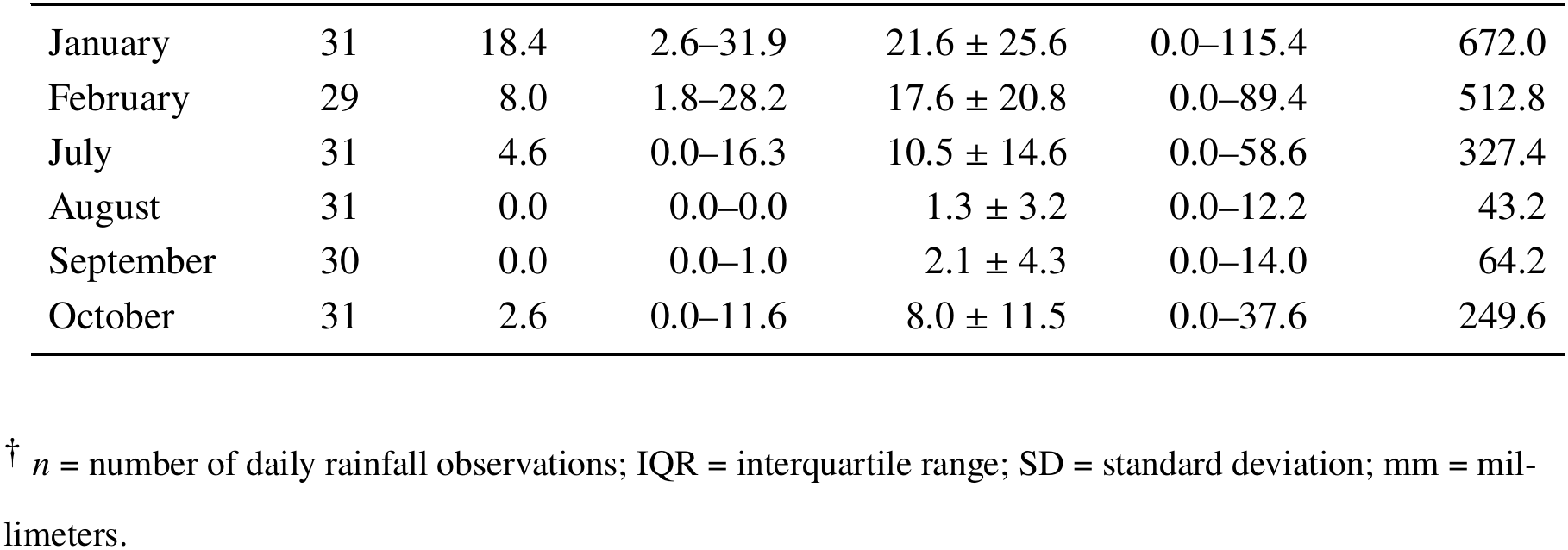
Daily rainfall (mm) at Ranomafana National Park, Madagascar, during sampled wet and dry season months in 2020.

**Figure 2.**
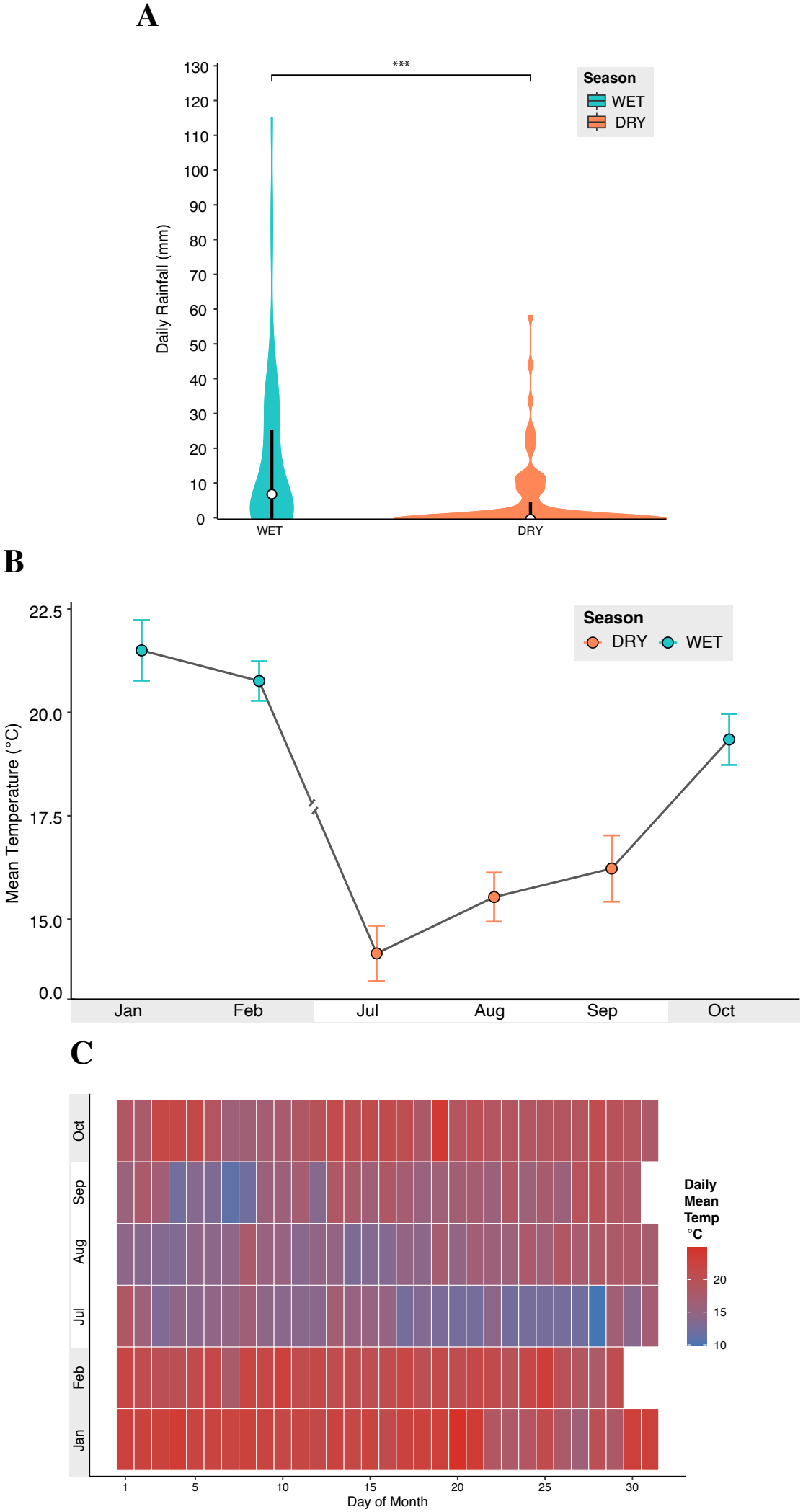
Significant differences in rainfall and temperature between wet and dry sampling periods at Ranomafana National Park, Madagascar, in 2020. **A)** Median daily rainfall by season. **B)** Mean monthly temperature with 95% confidence intervals. **C)** Mean daily air temperature across the study period.

Wet season temperatures averaged 20.53 ± 1.89 °C, which was significantly higher than 15.30 ± 2.04 °C during the dry season (Welch’s *t*(180.26) = 17.96, *p* = 5.03 × 10−42; Figure S2). Temperature also differed significantly among the six selected months (Kruskal-Wallis 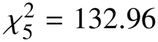, *p* = 5.59 × 10−27; Figures 2B and 2C). January had the highest median daily temperature, followed by February and October (Table 2). August was the coldest month. Dunn post-hoc tests showed a consistent seasonal pattern, with each wet season month differing significantly from each dry season month (all Holm-adjusted *p* < 0.001, Table S4). No pairwise comparison among the three wet season months remained significant after Holm adjustment, nor did any among the three dry season months.

**Table 2.**
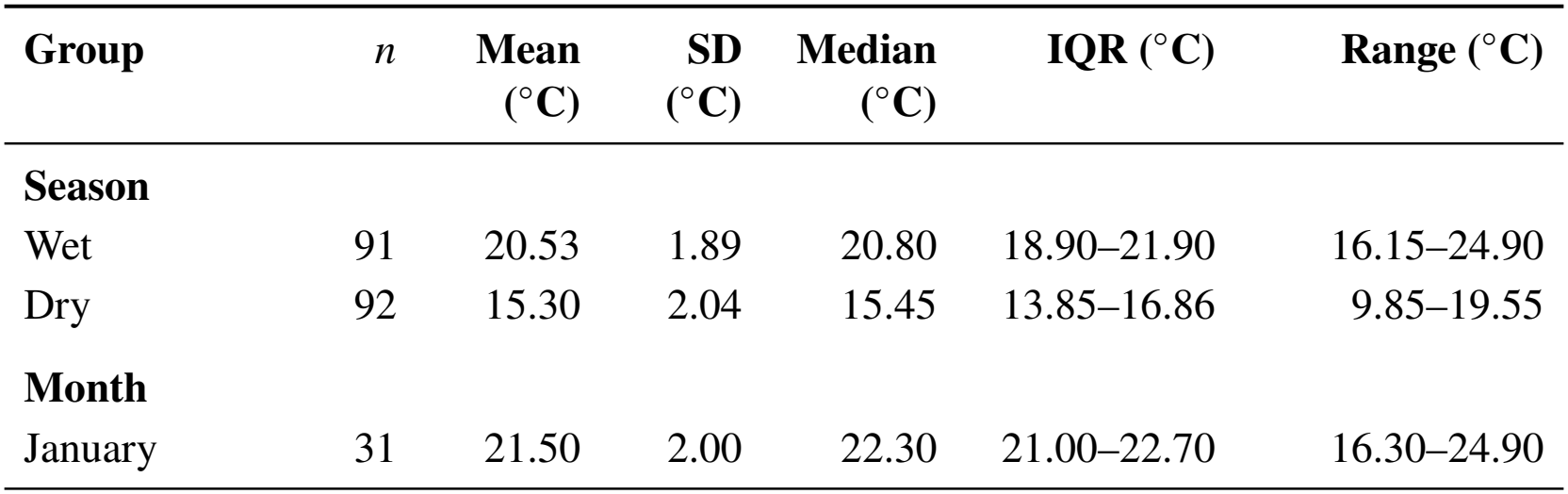
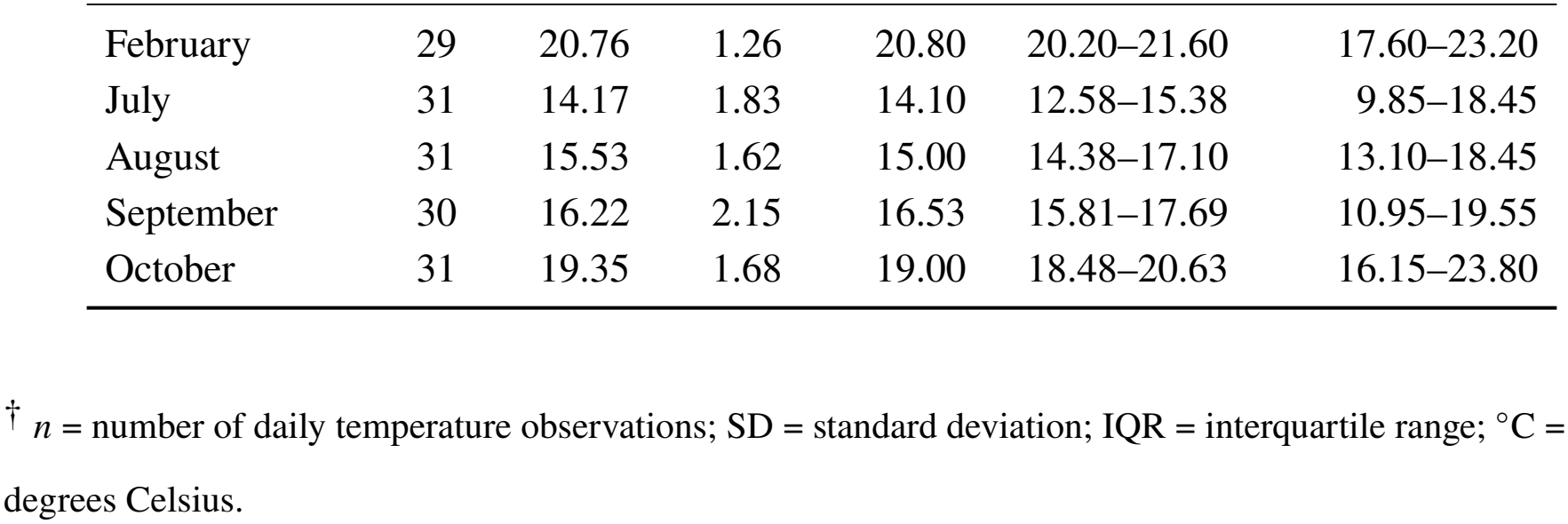
Daily temperature (°C) at Ranomafana National Park, Madagascar, during sampled wet and dry season months in 2020.

| **Group** | *n* | **Mean (°C)** | **SD (°C)** | **Median (°C)** | **IQR (°C)** | **Range (°C)** |
|---|---|---|---|---|---|---|
| **Season** |  |  |  |  |  |  |
| Wet | 91 | 20.53 | 1.89 | 20.80 | 18.90–21.90 | 16.15–24.90 |
| Dry | 92 | 15.30 | 2.04 | 15.45 | 13.85–16.86 | 9.85–19.55 |
| **Month** |  |  |  |  |  |  |
| January | 31 | 21.50 | 2.00 | 22.30 | 21.00–22.70 | 16.30–24.90 |
| February | 29 | 20.76 | 1.26 | 20.80 | 20.20–21.60 | 17.60–23.20 |
| July | 31 | 14.17 | 1.83 | 14.10 | 12.58–15.38 | 9.85–18.45 |
| August | 31 | 15.53 | 1.62 | 15.00 | 14.38–17.10 | 13.10–18.45 |
| September | 30 | 16.22 | 2.15 | 16.53 | 15.81–17.69 | 10.95–19.55 |
| October | 31 | 19.35 | 1.68 | 19.00 | 18.48–20.63 | 16.15–23.80 |

<sup>†</sup> *n* = number of daily temperature observations; SD = standard deviation; IQR = interquartile range; °C = degrees Celsius.

### 3.2 Diet and Behavior

The final dataset contained 3,278 behavioral observations across 30 focal-follow days, with 11 days during the wet and 19 during the dry season.

Multinomial models revealed significant overall differences in diet composition between seasons (Wald 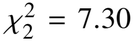, *p*= 0.026). Fruit consumption was observed for 87.57% of wet season observations while the dry season was marked by more varied diets, with increased consumption of leaves (18.13%) and other fallback foods (21.24%) (Figure 3A and Figure S3). The monthly diet model found overall differences in diet over the sampling period (Wald 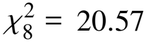, *p* = 0.008), but only one pairwise test (January – September) remained significant after Holm corrections (Wald 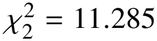, *p*= 0.035; Tables S5 and S6). Frugivory was highest in February (94.03%), folivory was highest in August (25.23%), and other fallback foods (i.e., flowers, seeds, and soil) were most consumed during September (36.17%) (Figure S4).

**Figure 3.**
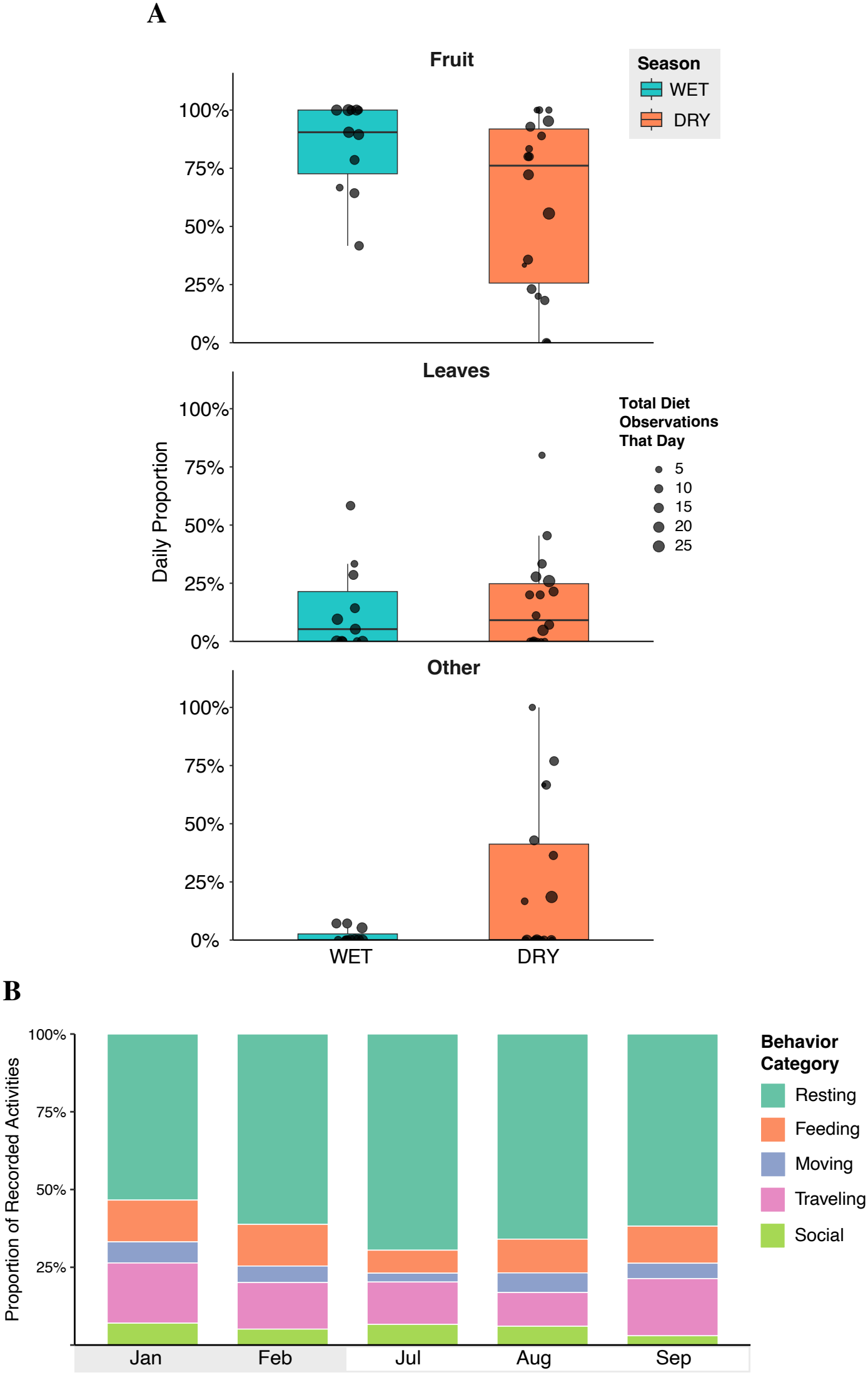
Seasonal and monthly variation in the diet and activity budgets of eastern *Eulemur rufifrons*. **A)** Daily proportions of consumed food categories observed between seasons (excluding October, data unavailable). Individual points represent sampling days while point size indicates the number of feeding observations recorded that day. Overall food category composition differed between seasons (multinomial mixed-effects logistic regression; joint Wald test, *p* < 0.05). **B)** Monthly activity budgets expressed as proportions of total behavioral observations each month.

Overall activity budget composition differed significantly among months, but pairwise tests did not provide enough evidence to identify specific months driving this pattern (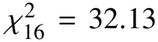, *p* = 0.0096, Tables S7 and S8 and Figure 3B). Overall activity budgets were not significantly different between seasons either (Wald 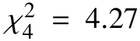, *p* = 0.371). Resting accounted for 56.4% of wet season observations and 66.1% of dry season observations, while feeding accounted for 13.4% and 10.1%, respectively (Table S9 and Figure S5).

### 3.3 Gut Microbiome Diversity, Composition, and Function

We generated 4,865,367 total sequence reads (min: 14,442; max: 404,579; mean: 79,760). We identified 2,606 unique bacterial ASVs before bioinformatic filtering. 3,197,112 sequence reads (min: 9,715; max: 269,733; mean: 52,411 per sample) were retained after quality filtering and rarefaction. 99.7% of sequence reads were assigned to known taxa.

We found cyclical fluctuations in the relative abundance of bacterial taxa over time, with samples from the beginning and end of our sampling period (both being wet seasons) harboring similar abundances (Figure 4A). There were more phyla present during the dry season, and more taxa categorized as “low abundance” (Table S10 and Figure S6).

**Figure 4.**
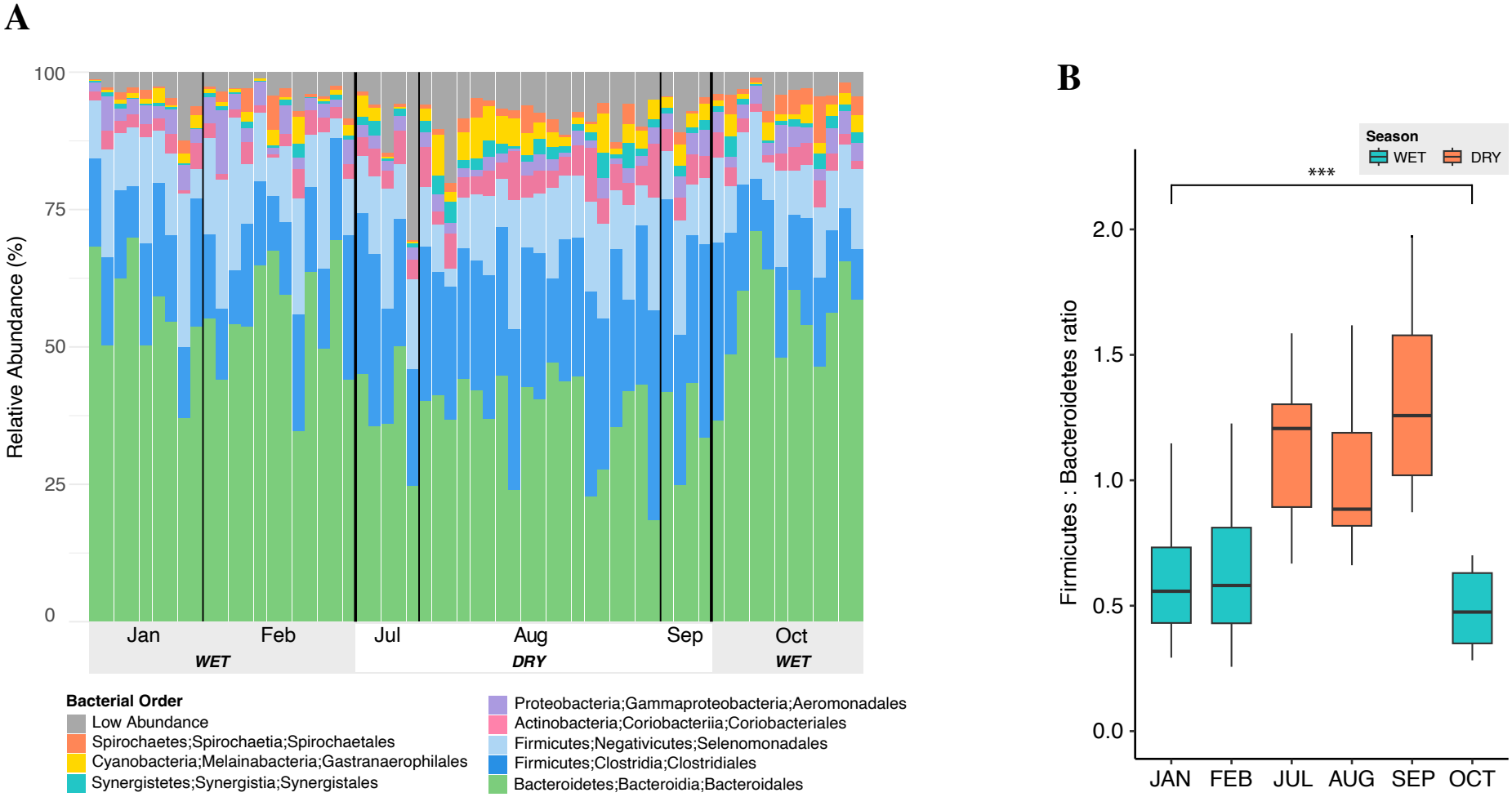
Variation in gut bacterial communities across sampling months. **A)** Relative abundance of bacterial taxa across individual fecal samples, grouped by month and season. **B)** Firmicutes-to-Bacteroidetes relative abundance ratios by month.

There was a seasonal trade-off in the Firmicutes-to-Bacteroidetes (F/B) ratio. F/B ratios differed significantly among sampling months (Kruskal–Wallis test: 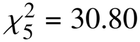, *p*< 0.001, Table S11 and Figure 4B) and between seasons (Kruskal–Wallis test: 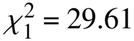, *p*< 0.001). The dry season had a higher median F/B ratio (median = 0.920, IQR = 0.821–1.34) than the wet season (median = 0.547, IQR = 0.408–0.701) (Figure S7).

Season and climate had significant effects on microbial richness and evenness (Figure 5). All three measures of alpha diversity were significantly higher during dry season months (Table 3, Figure 5A, and Figure S8). Kruskal-Wallis tests for each metric found season to have a stronger effect than month, although both were significant. There was no significant effect of sex nor individual identity on alpha diversity (Table 3).

**Table 3.** Significant associations of sampling season and month with gut microbiome alpha diversity in *Eulemur rufifrons* from Kruskal–Wallis tests.

| **Alpha Diversity Metric** | **Study Variable** | *H* | *p* |
|---|---|---|---|
| Faith's PD | Season | 20.1298 | < 0.0001<sup>***</sup> |
|  | Month | 21.1144 | 0.0007<sup>***</sup> |
|  | Sex | 0.1198 | 0.7291 |
|  | Individual | 7.3410 | 0.1964 |
| Shannon | Season | 35.7287 | < 0.0001<sup>***</sup> |
|  | Month | 36.3911 | < 0.0001<sup>***</sup> |
|  | Sex | 2.2509 | 0.1335 |
|  | Individual | 5.1979 | 0.3922 |
| Observed OTUs | Season | 13.0415 | 0.0003<sup>***</sup> |
|  | Month | 14.8974 | 0.0108<sup>*</sup> |
|  | Sex | 1.0054 | 0.3160 |
|  | Individual | 8.4336 | 0.1338 |

<sup>†</sup> Faith's PD = Faith's phylogenetic diversity; OTUs = operational taxonomic units; *H* = Kruskal–Wallis test statistic. *p*-values are shown with significance indicators: <sup>\*</sup>*p* < 0.05, <sup>\*\*</sup>*p* < 0.01, and <sup>\*\*\*</sup>*p* < 0.001.

**Figure 5.**
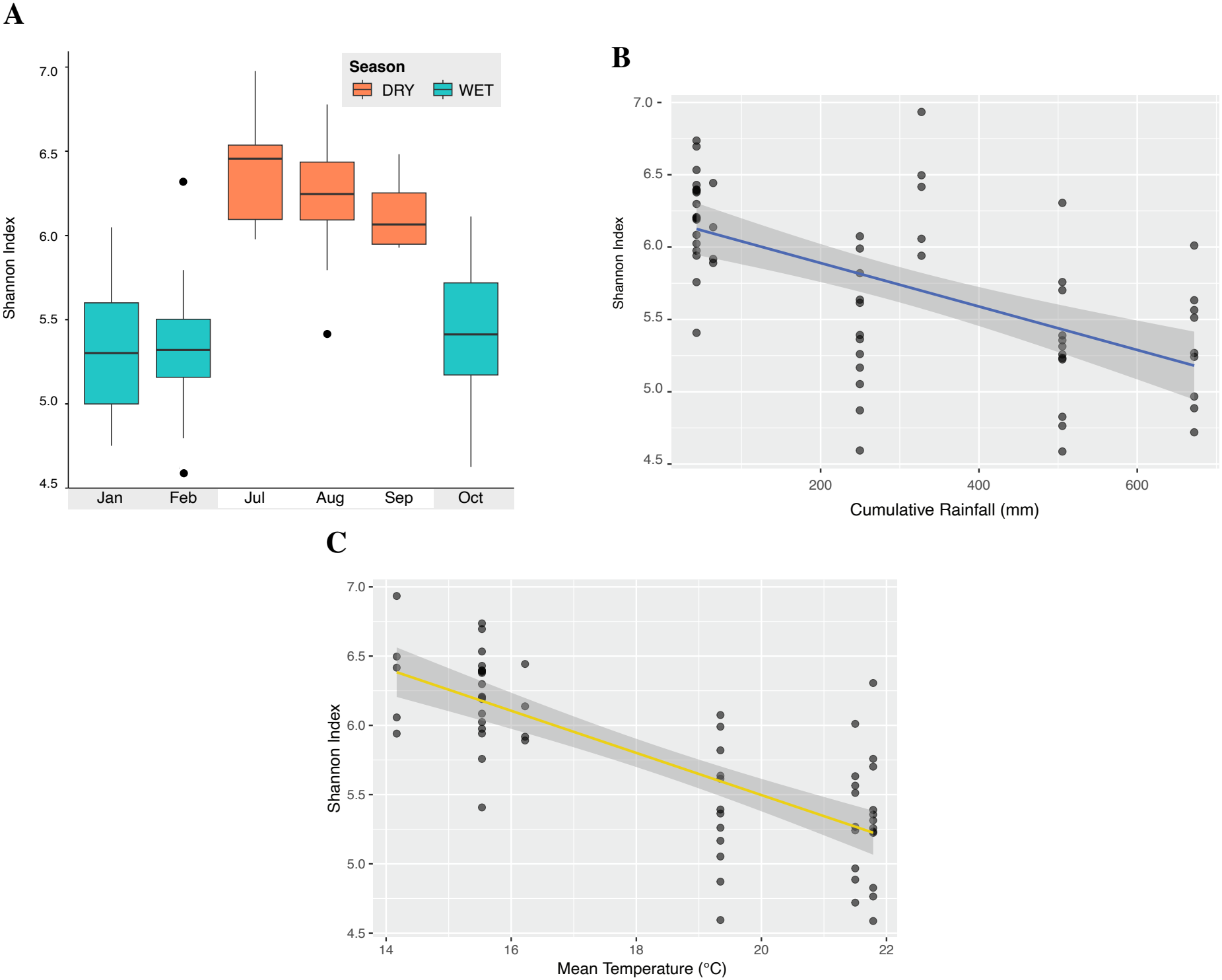
Significant associations of gut microbiome alpha diversity with sampling month and climate. **A)** Monthly Shannon diversity index by sampling season, with higher diversity during dry months (orange). **B)** Negative linear associations between Shannon Index and monthly cumulative rainfall. **C)** Negative linear associations between Shannon Index and monthly mean temperature.

LMMs for all three metrics of alpha diversity found moderate but significant negative associations between rainfall, temperature, and GM diversity with small random intercept variance (*p <* 0.001 for all) (Figure S9). Of the three metrics, LMMs for Shannon Index best met model assumptions and had the most normally distributed residuals (Figures 5B and 5C).

PCoA plots of all four beta diversity metrics showed samples clustering by season in ordination space (Figure S10). Adonis tests of all four beta diversity metrics found season to be the strongest explanatory variable (*p* = 0.001 for all metrics) (Table 4). Bray–Curtis dissimilarity (Figure 6A) and Weighted UniFrac plots exhibited the clearest clustering by season. Sex was only a significant variable for one metric (Bray–Curtis dissimilarity).

**Table 4.** Significant associations of sampling season and month with gut microbiome beta diversity in *Eulemur rufifrons* from PERMANOVA tests.

| **Beta Diversity Metric** | **Study Variable** | *R*<sup>2</sup> | *p* |
|---|---|---|---|
| Bray–Curtis | Season | 0.1773 | 0.001<sup>**</sup> |
|  | Month | 0.0842 | 0.004<sup>**</sup> |
|  | Season:Month | 0.0842 | 0.007<sup>**</sup> |
|  | Sex | 0.0229 | 0.033<sup>*</sup> |
|  | Individual | 0.0296 | 0.241 |
| Jaccard | Season | 0.0594 | 0.001<sup>**</sup> |
|  | Month | 0.0745 | 0.020<sup>*</sup> |
|  | Season:Month | 0.0745 | 0.038<sup>*</sup> |
|  | Sex | 0.0162 | 0.295 |
|  | Individual | 0.0352 | 0.144 |
| Weighted UniFrac | Season | 0.3630 | 0.001<sup>**</sup> |
|  | Month | 0.0513 | 0.228 |
|  | Season:Month | 0.0513 | 0.241 |
|  | Sex | 0.0200 | 0.091 |
|  | Individual | 0.0209 | 0.362 |
| Unweighted UniFrac | Season | 0.0929 | 0.001<sup>**</sup> |
|  | Month | 0.0757 | 0.080 |
|  | Season:Month | 0.0757 | 0.088 |
|  | Sex | 0.0113 | 0.782 |
|  | Individual | 0.0341 | 0.214 |

<sup>†</sup> PERMANOVA = permutational multivariate analysis of variance; UniFrac = unique fraction distance; *R*<sup>2</sup> = proportion of variation explained by the study variable. *p*-values are shown with significance indicators: <sup>\*</sup>*p* < 0.05, <sup>\*\*</sup>*p* < 0.01, and <sup>\*\*\*</sup>*p* < 0.001.

**Figure 6.**
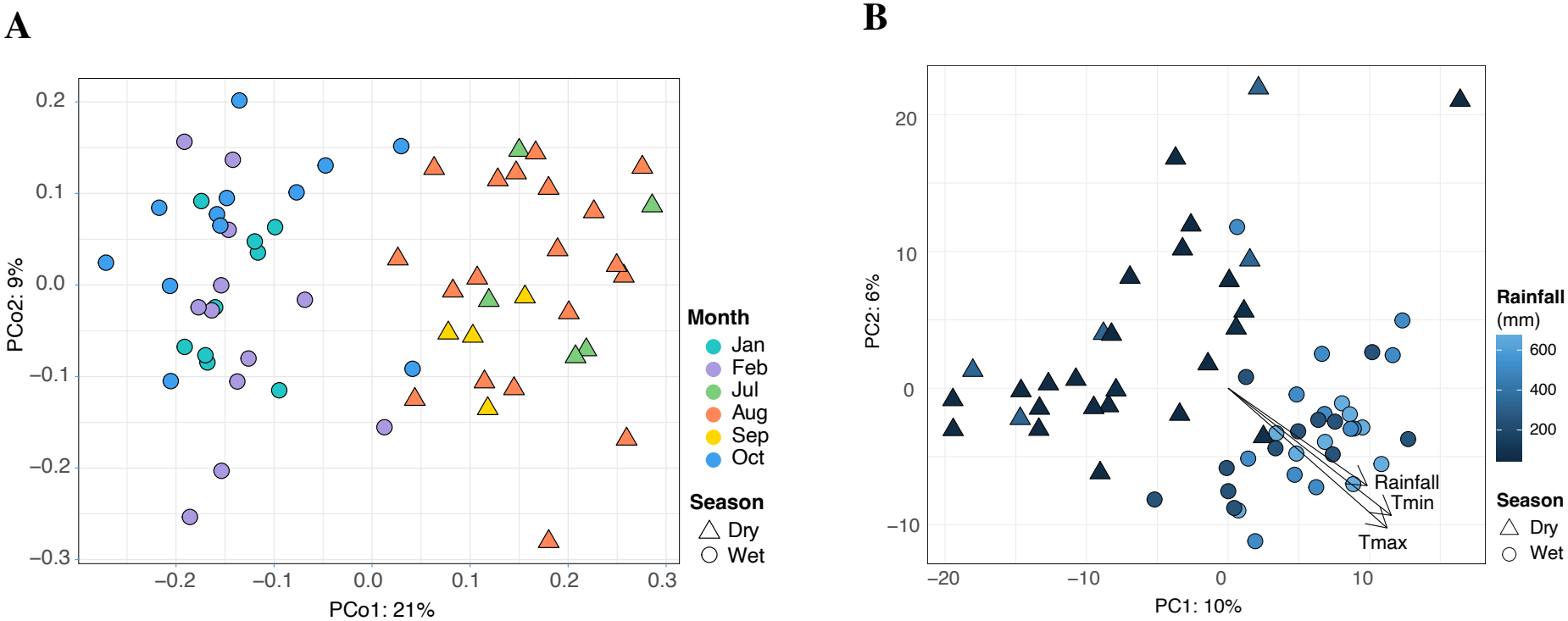
Seasonal and climatic patterns in gut microbiome beta diversity. **A)** Principal coordinates analysis (PCoA) of Bray–Curtis dissimilarity showing samples clustered by month (colors) and season (shapes). Samples are clustered by wet (circles) and dry (triangles) season in ordination space. **B)** Ordination of gut microbiome composition in relation to monthly rainfall and temperature; environmental vectors indicate the direction of association.

Climate-related environmental vectors exhibited directional influence on PCA ordination, suggesting that rainfall (R^2^ = 0.47, *p* = 0.001), minimum temperature (*R*^2^ = 0.73, *p* = 0.001), and maximum temperature (*R*^2^ = 0.70, *p* = 0.001) were major drivers of variation (Figure 6B). Similar to beta diversity plots, wet and dry season samples formed distinct clusters. Sex was not a significant variable (*R*^2^ = 0.002, *p* = 0.89).

The random forest classifier correctly predicted the season for 84.6% of our samples. Of true dry season samples, only 22.2% were misclassified as wet season. Of true wet season samples, 100% were correctly classified. 40 of the 50 most important taxa for distinguishing between wet and dry season samples belonged to the phyla *Firmicutes* and *Bacteroidetes* (Table S12 and Figure S11).

PICRUSt2 revealed seasonal differences in the predicted metabolic potential of gut microbial communities. Wilcoxon rank-sum tests identified 120 MetaCyc pathways that differed significantly between wet and dry season samples, with 62 pathways exhibiting greater predicted abundances during the wet season and 58 during the dry (all *p* < 0.0001; Table S13 and Figure S12). Wet season samples showed greater predicted abundances of several lipid, cofactor, and nucleotide biosynthesis pathways. In contrast, dry season samples had a greater representation of amino-acid biosynthesis pathways. Some of the largest seasonal differences observed were enriched fatty-acid biosynthesis pathways during the wet season. Wet season samples exhibited greater predicted abundances of pathways associated with pantothenate, folate, riboflavin, cobalamin, and pyrimidine nucleotide biosynthesis. Dry season samples exhibited greater predicted abundances of several sugar-acid degradation pathways and amino-acid biosynthesis pathways, including those associated with arginine, histidine, lysine, and tryptophan biosynthesis (Figure 7). Overall, the analysis identified the most common metabolic pathways in the entire dataset as: anaerobic gondoate biosynthesis, the Calvin-Benson-Bassham cycle, the non-oxidative pentose phosphate pathway, cis-vaccenate biosynthesis, and superpathway of pyrimidine nucleobases salvage. The top 10 pathways were found in approximately equal abundances.

**Figure 7.**
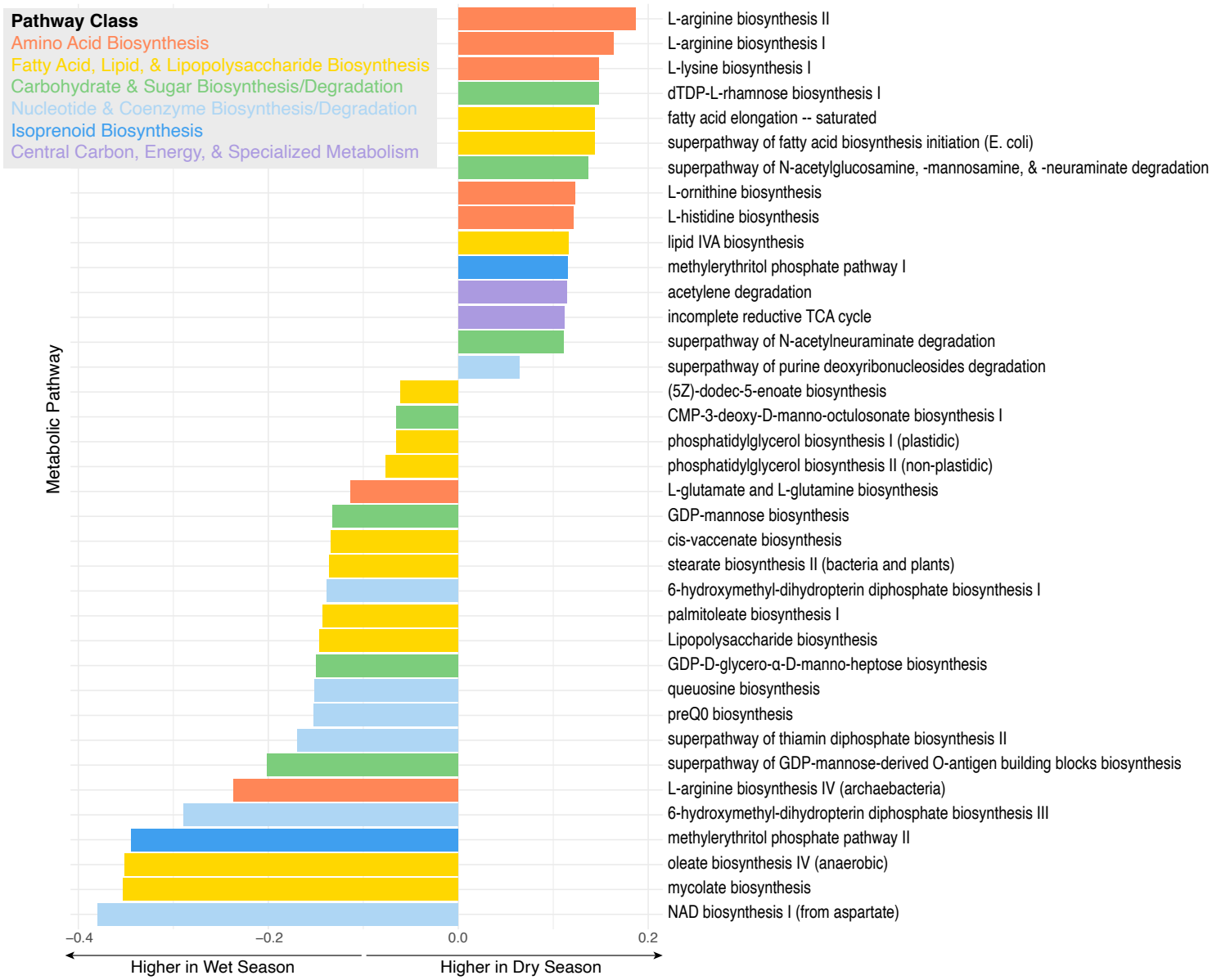
Seasonal differences in predicted metabolic pathways in the gut microbiome. Bars show pathways differing between wet and dry season samples, with colors distinguishing MetaCyc pathway classes. Only pathways with *p* < 1 × 10^-9^ are shown for optimal visualization.

## 4 Discussion

This study provides evidence for the relationship between seasonal shifts in climate, diet, and gut microbiota in a frugivorous, rainforest-dwelling primate. We expected eastern *E. rufifrons* to exhibit stronger seasonal shifts than previously detected in dry forest populations due to their habitat’s more pronounced seasonality. While inter-population comparisons were not possible, we observed characteristic wet and dry season profiles in terms of GM composition, microbial diversity, and predicted function over a short sampling period. Our findings support previously described cyclical rhythms, showing how GMs can revert to previous states when encountering familiar conditions (Beeby et al., 2025; Fan et al., 2022; Murillo, Schneider, Fichtel, and Daniel, 2022). Our data suggest that this 10-month study captured one full GM cycle, as samples from the beginning of the sampling period were compositionally similar to those at the end, despite a significant shift during the dry season in between (Figure 6A). These seasonal shifts demonstrate the capacity for rapid GM plasticity in response to ecological change, which may serve to enhance host metabolic flexibility during nutritional challenges.

Synchrony between changes in diet and GM diversity, composition, and function point to seasonal energy and nutrient discrepancies as the main driver of GM seasonality. First, our results suggest a positive relationship between dietary and GM diversity, as alpha diversity was higher when a wider range of fallback foods were consumed during the dry season (Figures S4 and S8). Additionally, dry season GMs harbored more taxa in the low abundance category (Figure S6). Greater presence of low abundance microbiota does not necessarily indicate changes in any resident gut services, as these taxa could be transient, acquired horizontally from a stochastic diet. According to the Anna-Karenina principle, such taxa may be temporary if they cannot successfully compete with established microbes for niche space (Zaneveld et al., 2017). However, we observed major seasonal shifts in the most abundant GM phyla, providing strong evidence that changes in gut function help compensate for dietary deficiencies. Fluctuating F/B ratios were the most prominent temporal changes observed, which are likely plastic responses to the need to digest carbohydrate-rich fruits versus protein-rich leaves. F/B ratios are a major GM benchmark due to the influence of these phyla on gut homeostasis and metabolite production (Y. Sun et al., 2023), and are highly sensitive to dietary treatments in both wild and experimental studies (Clayton et al., 2018; Xia et al., 2021). We found that gut Bacteroidetes were a hallmark of wet season frugivory. Bacteroidetes help initiate digestion of complex carbohydrates and non-structural polysaccharides into simple sugars, which can be utilized by the host and facilitate the proliferation of other gut bacteria (Zafar and Saier, 2021). In geladas, Bacteroidetes increase when hosts consume starchy roots and tubers, while Firmicutes increase when they consume a grass-heavy diet (Baniel et al., 2021). The GMs of frugivorous lemurs are typically dominated by microbiota specializing in slow fermentation of non-structural polysaccharides versus plant fiber in florivorous species (Greene et al., 2021). Prevotella species, some of the most abundant commensal gut bacteria in mammalian hosts (Tett et al., 2021), were largely responsible for the increase in Bacteroidetes abundance during the wet season (Figure 4A). Prevotella are a prominent gut fermenter of complex polysaccharides and a signature of fruit consumption in many primates, increasing in abundance during rainy seasons (Baniel et al., 2021; Xia et al., 2021). However, high Bacteroidetes abundance relative to other phyla between both seasons confirms that frugivorous primates still require specialized carbohydrate degradation year-round.

Resorting to folivory during fruit shortages may present nutritional challenges, requiring gut microbiota to facilitate high-fiber metabolism. While *E. rufifrons* defaulted to energy-rich fruit during the wet season, we observed increased supplementation with leaves and flowers during the dry season, which may provide higher protein and mineral content (Irwin et al., 2025). When modeling these effects on GM function, we identified more amino acid-associated gene pathways during the dry season (Figures 7 and S12). Clostridiales, especially *Lachnospiraceae*, were largely responsible for the higher proportion of Firmicutes present during the dry season. *Lachnospiraceae* produce short-chain fatty acids to extract energy from hemicellulose, cellulose, and lignin fibers. They are abundant in species that consume high-fiber diets, but have been positively correlated with tough leaf consumption in folivorous lemurs (e.g., *Propithecus verreauxi*, Greene et al., 2025; Springer et al., 2017), frugivorous lemurs (*Varecia variegata*, Beeby and Baden, 2021), and frugo-folivorous lemurs that consume highly seasonal diets (*Propithecus coquereli*, Greene et al., 2021). These taxa and others in the Firmicutes phylum (e.g., *Ruminococcaceae*) are maintained by daily dietary fiber and perform essential gut digestive functions, namely depolymerization of non-digestible oligosaccharides and resistant starches into healthy metabolites like butyric and lactic acid (Y. Sun et al., 2023). Overall, shifts in GM function likely serve to maximize energy and nutrients during the fruit-lean season. A recent study of frugivorous *Varecia variegata* found similar seasonal GM patterns, but nutrient intake was even more predictive of microbial abundances than dietary breakdown alone (Beeby et al., 2025). Thus, nutrient and caloric intake are likely the largest pressures driving GM shifts during times of preferred food scarcity.

Aside from diet, subtle seasonal differences in lemur behavior could have been reflected in the GM. Resting time peaked during the coldest months (July and August), which is expected given lemurs increase thermoregulatory behaviors as fruit availability and feeding time decrease (Donati et al., 2011). *E. rufifrons* also spent less time feeding during these months, consistent with previous observations of *Eulemur* species adopting energy-minimizing feeding and foraging strategies during food shortages (Donati et al., 2011; Mandl et al., 2018; Sato et al., 2016). Lemurs traveled less during the dry season, which is best explained as a lean-season energy conservation strategy observed in other lemurs (Mandl et al., 2018). Other *Eulemur* species, like *E. fulvus*, have been observed to travel less but feed more to maximize energy during the lean season (Rahalinarivo et al., 2023). However, *E. rufifrons* have previously been observed to expand, not restrict, their feeding range (known as habitat shifting) during times of fruit scarcity (Overdorff, 1993) and to increase proximity to water sources (Amoroso et al., 2020). Seasonal habitat shifting may increase GM diversity via environmental acquisition of microbes, such as from soil, which vary by geography, levels of habitat degradation, human contact, and other context-dependent factors (Bornbusch et al., 2022). These microbes may subsequently colonize and proliferate in the gut, though resident microbes influenced by diet generally out-compete transient taxa. Furthermore, microbial transmission between individuals may vary with seasonal behavior changes. Lemurs have been observed to increase mate grooming and huddling during cooler seasons as they rest more and feed less (Donati et al., 2011). Social group-specific seasonal shifts in GM composition and alpha diversity have been described in *Propithecus verreauxi* (Perofsky et al., 2021) and *Eulemur rubriventer*, including negative correlations between individual sociality and GM alpha diversity (Raulo et al., 2018).

The timing of annual rainfall and temperature changes were correlated with GM shifts, though it is difficult to disentangle the relative influence of environmental variables on the GM. Both rainfall and temperature were negatively correlated with alpha diversity and exhibited a directional influence on beta diversity (Figure 5 and Figure 6B). Previously, rainfall has overshadowed temperature as the stronger predictor of GM communities of some primates (including dry forest *E. rufifrons*; Murillo, Schneider, Heistermann, et al., 2022), likely linked to its influence on edible vegetation (Baniel et al., 2021; Beeby et al., 2025; Orkin et al., 2019). That said, higher temperatures have been linked to decreased *Firmicutes* and increased *Bacteroidetes* in short-faced moles (Xu et al., 2025), flying squirrels (Liu et al., 2019), and pigeons (Dietz et al., 2022). Climate–GM associations could reflect effects of rainfall and temperature on food availability and/or environmentally-acquired microbes. In either case, resemblances among alpha diversity, beta diversity, and rainfall patterns highlight the GM’s sensitivity to the host environment.

Climatic anomalies during our study period provided a unique opportunity to demonstrate real-time GM plasticity. Before analyzing climate data, we expected October GM profiles to resemble those from other dry season months, following the conventional seasonal calendar and previous *E. rufifrons* studies (Murillo, Schneider, Heistermann, et al., 2022). However, local climate records from RNP revealed an unusually early increase in rainfall and temperature during October, coinciding with GMs reverting to compositions observed in the earlier wet season months (Figure 4A). These changes occurred during a period of broader climate extremes across Madagascar in 2020, including unusually high land surface temperatures and the most severe drought seen in 30 years (Gouveia et al., 2024). Our findings suggest that the GM responds quickly to short-term climatic variation, a phenomenon which may become increasingly relevant as climate change continues to disrupt historical seasonal onsets and weather patterns. Climate change poses a mounting threat to lemurs in already-shrinking forest fragments by intensifying cyclones, exacerbating water demands, and increasing parasite and disease risk (Amoroso et al., 2020; Barrett et al., 2013; Rigden et al., 2024). Climate change-driven food shortages can be particularly threatening to host health, as fecal glucocorticoid data suggest that food scarcity may increase hypothalamic-pituitary-adrenal axis activation and possibly stress-related dysbiosis (Martínez-Mota et al., 2022; Murillo, Schneider, Heistermann, et al., 2022). However, whether climate-driven environmental changes disrupt GM dynamics and ultimately compromise host resiliency remains an important question for future studies.

With only 16S metabarcoding data and a limited sampling scope, we draw cautious conclusions about the speed, magnitude, and function of GM plasticity. However, we show that even imperfect sampling of a single population through time can uncover significant intra-species GM variation, even over short timescales. A major limitation of this study was a sampling gap between March and June due to the closure of RNP as a COVID-19 precaution. Nevertheless, the significant GM shift observed between September and October, coinciding with climatic and dietary changes, provides evidence for how rapidly the GM can respond to ecological change. Future longitudinal studies with repeated sampling across lifespans, ideally in tandem with ecological and dietary data, are needed to distinguish environmental pressures from drift. Metadata analyses, including evaluation of food nutritional content, could help parse out the relative contributions of diet, behavior, and climate on GM seasonality. Studies utilizing metagenomic and metatranscriptomic data can more expansively evaluate the functional contributions and metabolic potential of the GM in this context. Since *E. rufifrons* are widely distributed, a future study might directly compare samples from both eastern and western regions to 1) test whether climate extremes affect GM seasonality, and 2) test for broader effects of geography and host genetics between eastern and western *E. rufifrons*, which may be genetically and morphologically distinct enough to be considered unique species (Everson et al., 2025). Further investigation into GM plasticity as an adaptive mechanism is especially relevant for the conservation of endangered lemurs facing compounding pressures from habitat disturbance, bushmeat hunting, and climate change.

## 5 Conclusions

This study demonstrates a cascading relationship between climate, host diet, and the GM that is indicative of real-time plastic responses to changing environmental pressures. GM seasonality likely serves to mediate dietary shifts, helping hosts compensate for nutritional deficits when preferred food resources are more scarce. We hypothesize that seasonal differences in the GM will become more apparent with increasingly extreme seasons. Future investigation into how disruptions to GM cycling may affect host health would offer insights into the gravity of climate change on endangered primates and, more broadly, how the GM confers host fitness in rapidly changing environments.

## 6 Acknowledgements

Thank you to Stony Brook University’s Institute for the Conservation of Tropical Environments (ICTE) for providing climate data from RNP. This work would not be possible without the Malagasy Institute for the Conservation of Tropical Ecosystems (MICET), ICTE, Centre ValBio, Ranomafana National Park, Madagascar National Parks, and the Madagascar Ministry of Environment, Forests, and Ecology.

## 7 Data Availability Statement

Sequence data are available under BioProject ID PRJNA1563099 at https://www.ncbi.nlm.nih.gov/bioproject/PRJNA1563099.

## Notes

https://www.ncbi.nlm.nih.gov/bioproject/PRJNA1563099

### Competing Interest Statement

The authors have declared no competing interest.

