## Supplementary_Information for "Synchronized Seasonal Shifts in the Climate, Diet, and Gut Microbiome of a Rainforest Lemur (*Eulemur rufifrons*)"

### Supporting Tables

**Table S1.** Sample IDs with the month and season of collection from Ranomafana National Park in 2020. Wet season samples ($n$ = 33) were collected January 22 – February 13 and October 1 – 22. Dry season samples ($n$ = 28) were collected July 17 – September 30.

| Sample ID | Month | Season |
|---|---|---|
| RNP1 | JAN | WET |
| RNP2 | JAN | WET |
| RNP3 | JAN | WET |
| RNP4 | JAN | WET |
| RNP5 | JAN | WET |
| RNP6 | JAN | WET |
| RNP7 | JAN | WET |
| RNP8 | JAN | WET |
| RNP9 | JAN | WET |
| RNP12 | FEB | WET |
| RNP13 | FEB | WET |
| RNP14 | FEB | WET |
| RNP15 | FEB | WET |
| RNP16 | FEB | WET |
| RNP17 | FEB | WET |
| RNP19 | FEB | WET |
| RNP22 | FEB | WET |
| RNP23 | FEB | WET |
| RNP26 | FEB | WET |
| RNP27 | FEB | WET |
| RNP28 | FEB | WET |
| RNP31 | JULY | DRY |
| RNP33 | JULY | DRY |
| RNP35 | JULY | DRY |
| RNP38 | JULY | DRY |
| RNP42 | JULY | DRY |
| RNP44 | AUG | DRY |
| RNP45 | AUG | DRY |
| RNP46 | AUG | DRY |
| RNP50 | AUG | DRY |
| RNP51 | AUG | DRY |

*Continued on next page*

*Table S1 continued*

| Sample ID | Month | Season |
|---|---|---|
| RNP52 | AUG | DRY |
| RNP55 | AUG | DRY |
| RNP56 | AUG | DRY |
| RNP57 | AUG | DRY |
| RNP60 | AUG | DRY |
| RNP61 | AUG | DRY |
| RNP62 | AUG | DRY |
| RNP63 | AUG | DRY |
| RNP64 | AUG | DRY |
| RNP65 | AUG | DRY |
| RNP66 | AUG | DRY |
| RNP67 | AUG | DRY |
| RNP68 | AUG | DRY |
| RNP70 | AUG | DRY |
| RNP73 | SEPT | DRY |
| RNP77 | SEPT | DRY |
| RNP80 | SEPT | DRY |
| RNP81 | SEPT | DRY |
| RNP83 | OCT | WET |
| RNP84 | OCT | WET |
| RNP86 | OCT | WET |
| RNP89 | OCT | WET |
| RNP90 | OCT | WET |
| RNP92 | OCT | WET |
| RNP94 | OCT | WET |
| RNP96 | OCT | WET |
| RNP97 | OCT | WET |
| RNP98 | OCT | WET |
| RNP99 | OCT | WET |
| RNP100 | OCT | WET |

**Table S2.** Daily cumulative rainfall and daily minimum/maximum temperatures associated with the collection date of each fecal sample. Climate data were collected at the Centre ValBio weather station in Ranomafana National Park, January–October 2020.

| Sample ID | Month | Season | Date | Rain (mm) | $T_{min}$ (°C) | $T_{max}$ (°C) |
|---|---|---|---|---|---|---|
| RNP1 | JAN | WET | 2020-01-22 | 24.4 | 17.8 | 19.7 |
| RNP100 | OCT | WET | 2020-10-22 | 0 | 14.1 | 23.5 |
| RNP12 | FEB | WET | 2020-02-04 | 1.8 | 16.7 | 23.1 |
| RNP13 | FEB | WET | 2020-02-06 | 9.8 | 18.3 | 24.9 |
| RNP14 | FEB | WET | 2020-02-06 | 9.8 | 18.3 | 24.9 |
| RNP15 | FEB | WET | 2020-02-07 | 0 | 17.5 | 17.7 |
| RNP16 | FEB | WET | 2020-02-07 | 0 | 17.5 | 17.7 |
| RNP17 | FEB | WET | 2020-02-07 | 0 | 17.5 | 17.7 |
| RNP19 | FEB | WET | 2020-02-07 | 0 | 17.5 | 17.7 |
| RNP2 | JAN | WET | 2020-01-23 | 34 | 17.1 | 18.9 |
| RNP22 | FEB | WET | 2020-02-10 | 0 | 19.4 | 26.5 |
| RNP23 | FEB | WET | 2020-02-10 | 0 | 19.4 | 26.5 |
| RNP26 | FEB | WET | 2020-02-12 | 0 | 16.8 | 25.8 |
| RNP27 | FEB | WET | 2020-02-12 | 0 | 16.8 | 25.8 |
| RNP28 | FEB | WET | 2020-02-13 | 7.2 | 15.8 | 25.8 |
| RNP3 | JAN | WET | 2020-01-24 | 17.6 | 17.1 | 20.6 |
| RNP31 | JULY | DRY | 2020-07-17 | 7.6 | 10.8 | 14.4 |
| RNP33 | JULY | DRY | 2020-07-17 | 7.6 | 10.8 | 14.4 |
| RNP35 | JULY | DRY | 2020-07-20 | 0 | 10.4 | 14.6 |
| RNP38 | JULY | DRY | 2020-07-21 | 1.4 | 9.4 | 15.6 |
| RNP4 | JAN | WET | 2020-01-27 | 1.2 | 15.4 | 17.2 |
| RNP42 | JULY | DRY | 2020-07-27 | 0 | 8.4 | 15.8 |
| RNP44 | AUG | DRY | 2020-08-04 | 2 | 10.9 | 15.7 |
| RNP45 | AUG | DRY | 2020-08-04 | 2 | 10.9 | 15.7 |
| RNP46 | AUG | DRY | 2020-08-07 | 0 | 10.2 | 18.9 |
| RNP5 | JAN | WET | 2020-01-28 | 0 | 15.5 | 24.3 |
| RNP50 | AUG | DRY | 2020-08-17 | 0 | 9.9 | 20 |
| RNP51 | AUG | DRY | 2020-08-17 | 0 | 9.9 | 20 |
| RNP52 | AUG | DRY | 2020-08-17 | 0 | 9.9 | 20 |
| RNP55 | AUG | DRY | 2020-08-18 | 0 | 8.3 | 20.5 |
| RNP56 | AUG | DRY | 2020-08-18 | 0 | 8.3 | 20.5 |
| RNP57 | AUG | DRY | 2020-08-19 | 0 | 13.6 | 20.6 |
| RNP6 | JAN | WET | 2020-01-29 | 2.4 | 15.7 | 20.5 |
| RNP60 | AUG | DRY | 2020-08-20 | 0 | 8.3 | 21.4 |
| RNP61 | AUG | DRY | 2020-08-20 | 0 | 8.3 | 21.4 |
| RNP62 | AUG | DRY | 2020-08-21 | 0 | 10.4 | 22.5 |
| RNP63 | AUG | DRY | 2020-08-21 | 0 | 10.4 | 22.5 |
| RNP64 | AUG | DRY | 2020-08-21 | 0 | 10.4 | 22.5 |
| RNP65 | AUG | DRY | 2020-08-26 | 0 | 15.1 | 21.8 |

*Continued on next page*

*Table S2 continued*

| Sample ID | Month | Season | Date | Rain (mm) | $T_{min}$ (°C) | $T_{max}$ (°C) |
|---|---|---|---|---|---|---|
| RNP66 | AUG | DRY | 2020-08-26 | 0 | 15.1 | 21.8 |
| RNP67 | AUG | DRY | 2020-08-26 | 0 | 15.1 | 21.8 |
| RNP68 | AUG | DRY | 2020-08-27 | 0 | 12.5 | 21.9 |
| RNP7 | JAN | WET | 2020-01-29 | 2.4 | 15.7 | 20.5 |
| RNP70 | AUG | DRY | 2020-08-27 | 0 | 12.5 | 21.9 |
| RNP73 | SEPT | DRY | 2020-09-04 | 0 | 9.5 | 14.8 |
| RNP77 | SEPT | DRY | 2020-09-22 | 0 | 9.4 | 22.5 |
| RNP8 | JAN | WET | 2020-01-30 | 4.4 | 20.3 | 25.5 |
| RNP80 | SEPT | DRY | 2020-09-29 | 0 | 13.2 | 23.6 |
| RNP81 | SEPT | DRY | 2020-09-30 | 0 | 12.1 | 24 |
| RNP83 | OCT | WET | 2020-10-01 | 0 | 12.1 | 25.1 |
| RNP84 | OCT | WET | 2020-10-12 | 0 | 15.7 | 23.5 |
| RNP86 | OCT | WET | 2020-10-13 | 7.2 | 17.4 | 24.3 |
| RNP89 | OCT | WET | 2020-10-14 | 2.6 | 17.5 | 23.2 |
| RNP9 | JAN | WET | 2020-01-30 | 4.4 | 20.3 | 25.5 |
| RNP90 | OCT | WET | 2020-10-19 | 35.2 | 13.2 | 34.4 |
| RNP92 | OCT | WET | 2020-10-20 | 6.2 | 16 | 22.1 |
| RNP94 | OCT | WET | 2020-10-20 | 6.2 | 16 | 22.1 |
| RNP96 | OCT | WET | 2020-10-21 | 0 | 16.6 | 23.4 |
| RNP97 | OCT | WET | 2020-10-21 | 0 | 16.6 | 23.4 |
| RNP98 | OCT | WET | 2020-10-22 | 0 | 14.1 | 23.5 |
| RNP99 | OCT | WET | 2020-10-22 | 0 | 14.1 | 23.5 |

† mm = millimeters; $T_{min}$ and $T_{max}$ = recorded daily minimum and maximum temperatures; °C = degrees Celsius.

**Table S3.** Pairwise comparisons of daily rainfall among sampled months in 2020 (Dunn post-hoc tests following a Kruskal–Wallis test). Positive Z values indicate a higher mean rank in the first month listed, while negative values indicate such in the second month listed.

| Comparison | $Z$ | Holm-adjusted $p$ |
|---|---|---|
| January vs. February | 0.46 | 1.000 |
| January vs. July | 1.83 | 0.353 |
| January vs. August | 5.23 | $2.50 \times 10^{-6***}$ |
| January vs. September | 4.53 | $7.68 \times 10^{-5***}$ |
| January vs. October | 2.39 | 0.134 |
| February vs. July | 1.34 | 0.721 |
| February vs. August | 4.68 | $3.94 \times 10^{-5***}$ |

*Continued on next page*

*Table S3 continued*

| **Comparison** | $Z$ | **Holm-adjusted** $p$ |
|---|---|---|
| February vs. September | 4.00 | $7.70 \times 10^{-4***}$ |
| February vs. October | 1.89 | 0.353 |
| July vs. August | 3.40 | $0.00739^{**}$ |
| July vs. September | 2.71 | 0.0601 |
| July vs. October | 0.56 | 1.000 |
| August vs. September | −0.66 | 1.000 |
| August vs. October | −2.84 | $0.0449^{*}$ |
| September vs. October | −2.16 | 0.217 |

† $Z$ = standardized Dunn test statistic. Significance symbols refer to adjusted values: $^{*}p < 0.05$, $^{**}p < 0.01$, and $^{***}p < 0.001$.

**Table S4.** Pairwise comparisons of daily mean temperature among sampled months in 2020 (Dunn's post hoc tests following a Kruskal–Wallis test). Positive Z values indicate a higher mean rank for the first-listed month and negative values for the second-listed month.

| **Comparison** | $Z$ | **Unadjusted** $p$ | **Holm-adjusted** $p$ |
|---|---|---|---|
| January vs. February | 0.92 | 0.358 | 0.707 |
| January vs. July | 8.81 | $1.25 \times 10^{-18}$ | $1.88 \times 10^{-17***}$ |
| January vs. August | 7.41 | $1.31 \times 10^{-13}$ | $1.70 \times 10^{-12***}$ |
| January vs. September | 6.42 | $1.39 \times 10^{-10}$ | $1.67 \times 10^{-9***}$ |
| January vs. October | 2.59 | 0.00951 | 0.0571 |
| February vs. July | 7.74 | $9.72 \times 10^{-15}$ | $1.36 \times 10^{-13***}$ |
| February vs. August | 6.36 | $2.00 \times 10^{-10}$ | $2.20 \times 10^{-9***}$ |
| February vs. September | 5.40 | $6.72 \times 10^{-8}$ | $6.04 \times 10^{-7***}$ |
| February vs. October | 1.63 | 0.103 | 0.412 |
| July vs. August | −1.41 | 0.160 | 0.480 |
| July vs. September | −2.32 | 0.0203 | 0.101 |
| July vs. October | −6.22 | $5.07 \times 10^{-10}$ | $5.07 \times 10^{-9***}$ |
| August vs. September | −0.93 | 0.354 | 0.707 |
| August vs. October | −4.81 | $1.49 \times 10^{-6}$ | $1.20 \times 10^{-5***}$ |
| September vs. October | −3.84 | $1.21 \times 10^{-4}$ | $8.44 \times 10^{-4***}$ |

† $Z$ = standardized Dunn test statistic. Significance symbols correspond to Holm-adjusted values: $^{*}p < 0.05$, $^{**}p < 0.01$, and $^{***}p < 0.001$.

**Table S5.** Observation counts and within-month proportions of consumed food categories during focal follows of *Eulemur rufifrons* in 2020.

| Measure | JAN | FEB | JUL | AUG | SEP |
|---|---|---|---|---|---|
| $n_{days}$ | 7 | 4 | 5 | 8 | 5 |
| $n_{observations}$ | 102 | 67 | 39 | 107 | 47 |
| $n_{fruit}$ | 85 | 63 | 31 | 57 | 29 |
| $n_{leaves}$ | 15 | 3 | 7 | 27 | 1 |
| $n_{other}$ | 2 | 1 | 1 | 23 | 17 |
| $\%_{fruit}$ | 83.3 | 94 | 79.5 | 53.3 | 61.7 |
| $\%_{leaves}$ | 14.7 | 4.5 | 17.9 | 25.2 | 2.1 |
| $\%_{other}$ | 2.0 | 1.5 | 2.6 | 21.5 | 36.2 |

† $n_{days}$ = the number of observation days; $n_{observations}$ = the number of identified feeding records; $n_{fruit}$, $n_{leaves}$, and $n_{other}$ = counts within each food category, with ”other” including flowers, seeds, and soil.

**Table S6.** Pairwise differences comparisons of monthly dietary composition in *Eulemur rufifrons* using joint Wald tests from a multinomial mixed-effects logistic regression. Wald tests jointly compare the proportions of fruit, leaves, and other foods between months, with fruit as the model reference category. Adjusted $p$-values use the Holm correction across 10 comparisons.

| Comparison | Wald $\chi^2$ | df | Raw $p$ | Adjusted $p$ |
|---|---|---|---|---|
| Jan–Feb | 1.662 | 2 | 0.4357 | 1 |
| Jan–Jul | 0.236 | 2 | 0.8886 | 1 |
| Jan–Aug | 5.189 | 2 | 0.0747 | 0.373 |
| Jan–Sep | 11.285 | 2 | 0.0035 | 0.035* |
| Feb–Jul | 2.336 | 2 | 0.3110 | 1 |
| Feb–Aug | 7.126 | 2 | 0.0284 | 0.230 |
| Feb–Sep | 6.893 | 2 | 0.0319 | 0.230 |
| Jul–Aug | 2.205 | 2 | 0.3321 | 1 |
| Jul–Sep | 7.337 | 2 | 0.0255 | 0.230 |
| Aug–Sep | 5.847 | 2 | 0.0537 | 0.322 |

† df = degrees of freedom.

**Table S7.** Monthly activity budgets across focal-follow observation days of *Eulemur rufifrons* in 2020. Percentages are calculated within each month. Moving refers to movements within trees; traveling refers to movement between trees; social behaviors include grooming, playing, fighting, vocalizing, and tail wagging.

| **Month** | **Days** | **Records** | **Feeding** | **Moving** | **Traveling** | **Resting** | **Social** |
|---|---|---|---|---|---|---|---|
| Jan | 7 | 796 | 13.4% | 6.8% | 19.3% | 53.4% | 7.0% |
| Feb | 4 | 508 | 13.4% | 5.3% | 15.0% | 61.2% | 5.1% |
| Jul | 5 | 528 | 7.4% | 2.8% | 13.6% | 69.5% | 6.6% |
| Aug | 9 | 1,043 | 10.8% | 6.3% | 10.8% | 66.0% | 6.0% |
| Sep | 5 | 403 | 11.9% | 5.0% | 18.4% | 61.8% | 3.0% |

**Table S8.** Pairwise comparisons of monthly activity-budget composition based on joint Wald chi-square tests from a multinomial mixed-effects logistic regression model. The joint test compares feeding, moving, traveling, resting, and social behavior, with resting as the reference category. $p$-values have been adjusted across the 10 monthly comparisons using the Holm procedure.

| **Comparison** | **Wald $\chi^2$** | **df** | **Holm-adjusted $p$** |
|---|---|---|---|
| Jan vs. Feb | 2.500 | 4 | 1.000 |
| Jan vs. Jul | 11.470 | 4 | 0.174 |
| Jan vs. Aug | 7.632 | 4 | 0.530 |
| Jan vs. Sep | 5.860 | 4 | 0.839 |
| Feb vs. Jul | 9.760 | 4 | 0.268 |
| Feb vs. Aug | 4.579 | 4 | 1.000 |
| Feb vs. Sep | 3.536 | 4 | 1.000 |
| Jul vs. Aug | 10.204 | 4 | 0.260 |
| Jul vs. Sep | 14.843 | 4 | 0.050 |
| Aug vs. Sep | 13.894 | 4 | 0.069 |

† df = degrees of freedom.

**Table S9.** Seasonal activity budgets across focal-follow observation days of *Eulemur rufifrons* in 2020. Percentages are calculated within season. Moving refers to movements within trees; traveling refers to movement between trees; social behaviors include grooming, playing, fighting, vocalizing, and tail wagging. Wet season behavioral observations occurred in January and February while dry season behavioral observations were between July and September.

| Season | Days | Records | Feeding | Moving | Traveling | Resting | Social |
|---|---|---|---|---|---|---|---|
| Wet | 11 | 1,304 | 13.4% | 6.2% | 17.6% | 56.4% | 6.3% |
| Dry | 19 | 1,974 | 10.1% | 5.1% | 13.1% | 66.1% | 5.6% |

**Table S10.** Order-level relative abundance of gut bacteria in individual *Eulemur rufifrons* samples.

| Sample | Clostridiales | Bacteroidales | Spirochaetales | Gastranaerophilales | Selenomonadales | Aeromonadales | Coriobacteriales | Synergistales | Low Abundance |
|---|---|---|---|---|---|---|---|---|---|
| RNP1 | 16.12 | 68.29 | 0.13 | 0.24 | 10.56 | 1.69 | 1.49 | 0.17 | 1.31 |
| RNP2 | 16.14 | 50.24 | 0.38 | 0.57 | 19.64 | 6.26 | 3.32 | 0.63 | 2.81 |
| RNP3 | 16.08 | 62.45 | 1.50 | 0.68 | 10.39 | 2.38 | 2.14 | 0.81 | 3.58 |
| RNP4 | 9.44 | 69.94 | 1.17 | 0.89 | 10.58 | 2.85 | 2.32 | 0.13 | 2.68 |
| RNP5 | 18.64 | 50.33 | 1.68 | 0.99 | 19.21 | 3.59 | 2.23 | 0.19 | 3.14 |
| RNP6 | 20.67 | 59.19 | 0.39 | 2.73 | 9.52 | 2.30 | 2.17 | 0.47 | 2.56 |
| RNP7 | 15.88 | 54.58 | 1.32 | 0.42 | 14.74 | 4.57 | 3.47 | 0.30 | 4.71 |
| RNP8 | 13.05 | 36.98 | 2.43 | 1.88 | 27.94 | 4.64 | 0.49 | 0.16 | 12.41 |
| RNP9 | 23.44 | 53.65 | 1.64 | 2.32 | 5.23 | 2.64 | 4.71 | 0.14 | 6.23 |
| RNP12 | 15.22 | 55.22 | 0.47 | 0.77 | 17.64 | 4.73 | 2.67 | 0.69 | 2.58 |
| RNP13 | 13.09 | 43.97 | 2.02 | 0.71 | 23.35 | 11.76 | 1.02 | 0.66 | 3.43 |
| RNP14 | 9.77 | 54.16 | 0.28 | 0.53 | 27.86 | 3.01 | 1.37 | 0.15 | 2.88 |
| RNP15 | 18.79 | 53.68 | 4.21 | 1.93 | 10.93 | 2.51 | 3.90 | 1.09 | 2.97 |
| RNP16 | 15.36 | 64.86 | 0.12 | 0.48 | 12.42 | 4.33 | 1.33 | 0.07 | 1.04 |
| RNP17 | 9.95 | 67.55 | 6.22 | 2.83 | 6.97 | 1.20 | 0.57 | 0.41 | 4.30 |
| RNP19 | 13.35 | 59.43 | 1.70 | 0.43 | 13.81 | 5.31 | 2.11 | 1.01 | 2.85 |
| RNP22 | 21.26 | 34.72 | 1.05 | 4.94 | 21.03 | 2.13 | 5.30 | 2.49 | 7.08 |
| RNP23 | 15.57 | 63.57 | 0.64 | 0.03 | 9.54 | 2.27 | 4.40 | 0.01 | 3.96 |
| RNP26 | 14.74 | 49.59 | 0.44 | 0.35 | 24.75 | 2.67 | 2.58 | 0.43 | 4.45 |
| RNP27 | 18.51 | 69.48 | 0.82 | 0.90 | 3.65 | 1.34 | 2.06 | 0.80 | 2.46 |
| RNP28 | 26.25 | 44.07 | 1.21 | 2.02 | 10.23 | 4.56 | 2.62 | 0.63 | 8.41 |
| RNP31 | 29.38 | 45.00 | 0.74 | 3.97 | 10.34 | 2.28 | 3.40 | 1.40 | 3.50 |
| RNP33 | 31.41 | 35.60 | 0.66 | 2.36 | 14.00 | 2.48 | 4.98 | 2.62 | 5.89 |
| RNP35 | 21.01 | 36.01 | 0.88 | 0.23 | 21.90 | 1.83 | 3.16 | 0.36 | 14.62 |
| RNP38 | 23.20 | 50.12 | 0.59 | 0.39 | 10.03 | 2.63 | 5.99 | 1.31 | 5.73 |
| RNP42 | 21.35 | 24.64 | 0.35 | 0.36 | 16.32 | 2.35 | 3.52 | 0.62 | 30.49 |
| RNP44 | 28.13 | 40.11 | 0.81 | 2.19 | 10.84 | 2.81 | 7.22 | 1.98 | 5.91 |
| RNP45 | 22.45 | 41.26 | 1.13 | 7.41 | 8.58 | 3.30 | 2.31 | 3.27 | 10.29 |
| RNP46 | 24.23 | 36.79 | 1.75 | 1.83 | 3.31 | 1.89 | 6.35 | 3.76 | 20.09 |

*Continued on next page*

*Table S10 continued*

| Sample | Clostridiales | Bacteroidales | Spirochaetales | Gastranaerophilales | Selenomonadales | Aeromonadales | Coriobacteriales | Synergistales | Low Abundance |
|---|---|---|---|---|---|---|---|---|---|
| RNP50 | 23.87 | 44.19 | 2.56 | 4.96 | 9.13 | 1.72 | 3.71 | 1.44 | 8.42 |
| RNP51 | 23.74 | 42.07 | 3.68 | 7.35 | 11.97 | 1.33 | 3.23 | 1.94 | 4.69 |
| RNP52 | 26.23 | 36.88 | 0.96 | 6.38 | 14.38 | 2.69 | 4.48 | 2.83 | 5.16 |
| RNP55 | 27.05 | 44.78 | 1.29 | 5.19 | 8.78 | 1.69 | 2.89 | 1.65 | 6.68 |
| RNP56 | 29.28 | 23.99 | 1.41 | 5.16 | 23.48 | 0.49 | 8.85 | 0.39 | 6.95 |
| RNP57 | 25.51 | 42.69 | 5.18 | 3.92 | 8.99 | 2.72 | 3.91 | 1.07 | 6.01 |
| RNP60 | 26.65 | 40.51 | 1.91 | 2.99 | 10.72 | 3.06 | 4.15 | 2.68 | 7.32 |
| RNP61 | 15.43 | 47.14 | 2.40 | 2.81 | 16.40 | 2.05 | 3.08 | 2.20 | 8.49 |
| RNP62 | 26.02 | 43.65 | 0.49 | 1.64 | 11.53 | 0.83 | 3.38 | 1.06 | 11.40 |
| RNP63 | 25.32 | 44.65 | 0.99 | 1.21 | 11.27 | 2.66 | 5.49 | 1.13 | 7.28 |
| RNP64 | 37.40 | 22.77 | 0.40 | 1.44 | 16.31 | 1.40 | 9.74 | 1.48 | 9.05 |
| RNP65 | 27.56 | 27.69 | 2.14 | 7.13 | 17.22 | 3.72 | 4.53 | 4.50 | 5.50 |
| RNP66 | 32.56 | 35.34 | 1.22 | 1.04 | 10.65 | 1.15 | 4.62 | 0.70 | 12.73 |
| RNP67 | 16.63 | 42.00 | 3.61 | 3.54 | 17.14 | 2.73 | 4.00 | 4.56 | 5.79 |
| RNP68 | 29.02 | 43.12 | 0.92 | 3.18 | 7.27 | 1.88 | 3.46 | 1.32 | 9.84 |
| RNP70 | 38.25 | 18.51 | 0.08 | 3.55 | 20.45 | 3.10 | 9.73 | 1.41 | 4.91 |
| RNP73 | 35.12 | 41.79 | 0.31 | 1.95 | 8.74 | 2.83 | 3.97 | 0.97 | 4.31 |
| RNP77 | 27.47 | 24.79 | 2.34 | 3.89 | 20.75 | 3.98 | 4.07 | 1.82 | 10.89 |
| RNP80 | 26.99 | 43.39 | 0.55 | 2.55 | 9.14 | 3.93 | 5.47 | 0.98 | 6.99 |
| RNP81 | 35.21 | 33.50 | 1.14 | 2.95 | 12.10 | 4.78 | 3.97 | 1.75 | 4.59 |
| RNP83 | 32.39 | 36.55 | 1.29 | 1.02 | 15.58 | 3.84 | 4.50 | 0.92 | 3.92 |
| RNP84 | 22.16 | 48.66 | 3.62 | 4.04 | 8.51 | 2.01 | 3.31 | 3.62 | 4.07 |
| RNP86 | 19.28 | 60.27 | 0.96 | 0.91 | 9.58 | 2.47 | 2.51 | 0.92 | 3.10 |
| RNP89 | 9.50 | 71.11 | 1.02 | 0.26 | 12.19 | 3.34 | 1.47 | 0.20 | 0.91 |
| RNP90 | 12.68 | 64.08 | 2.18 | 3.14 | 6.82 | 2.32 | 1.31 | 0.36 | 7.11 |
| RNP92 | 16.61 | 48.00 | 3.73 | 1.15 | 17.48 | 5.32 | 2.99 | 0.70 | 4.03 |
| RNP94 | 13.82 | 60.31 | 4.62 | 0.78 | 8.06 | 3.79 | 4.20 | 1.30 | 3.12 |
| RNP96 | 19.59 | 53.91 | 2.98 | 3.40 | 9.73 | 2.91 | 3.91 | 0.62 | 2.96 |
| RNP97 | 16.33 | 46.34 | 8.50 | 2.04 | 12.77 | 2.14 | 4.85 | 2.70 | 4.34 |
| RNP98 | 15.07 | 56.26 | 1.71 | 2.21 | 10.77 | 4.58 | 3.11 | 2.11 | 4.18 |
| RNP99 | 9.72 | 65.59 | 2.08 | 2.07 | 11.50 | 4.27 | 1.87 | 1.11 | 1.79 |
| RNP100 | 9.21 | 58.64 | 3.50 | 2.98 | 14.52 | 3.33 | 1.47 | 1.91 | 4.42 |

**Table S12.** Top 50 bacterial features ranked by importance in a random forest classifier that predicts sampling season from gut microbiome composition. Larger importance values indicate greater contribution to discriminating between wet and dry season samples.

| Importance | Taxon |
|---|---|
| 0.026034901 | Bacteria;Firmicutes;Clostridia;Clostridiales;Lachnospiraceae |
| 0.023096692 | Bacteria;Firmicutes;Clostridia;Clostridiales;Ruminococcaceae |
| 0.02269274 | Bacteria;Firmicutes;Clostridia;Clostridiales;Lachnospiraceae |
| 0.022673718 | Bacteria;Bacteroidetes;Bacteroidia;Bacteroidales;Prevotellaceae |

*Continued on next page*

*Table S12 continued*

| Importance | Taxon |
|---:|---|
| 0.021065026 | Bacteria;Bacteroidetes;Bacteroidia;Bacteroidales;Prevotellaceae |
| 0.020765582 | Bacteria;Synergistetes;Synergistia;Synergistales;Synergistaceae;Pyramidobacter;uncultured bacterium |
| 0.019224074 | Bacteria;Bacteroidetes;Bacteroidia;Bacteroidales;Prevotellaceae |
| 0.018046653 | Bacteria;Firmicutes;Clostridia;Clostridiales;Lachnospiraceae |
| 0.017856131 | Bacteria;Bacteroidetes;Bacteroidia;Bacteroidales;Rikenellaceae;Rikenellaceae RC9 gut group |
| 0.017689062 | Bacteria;Bacteroidetes;Bacteroidia;Bacteroidales;Prevotellaceae;Alloprevotella;uncultured bacterium |
| 0.016513284 | Bacteria;Firmicutes;Negativicutes;Selenomonadales;Acidaminococcaceae;Phascolarctobacterium |
| 0.01609215 | Bacteria;Firmicutes;Clostridia;Clostridiales;Ruminococcaceae;Ruminococcaceae UCG-005;metagenome |
| 0.016038127 | Bacteria;Firmicutes;Clostridia;Clostridiales;Lachnospiraceae |
| 0.015446355 | Bacteria;Firmicutes;Clostridia;Clostridiales;Eubacteriaceae;Anaerofustis;uncultured bacterium |
| 0.01527014 | Bacteria;Proteobacteria;Deltaproteobacteria;Desulfovibrionales;Desulfovibrionaceae;Desulfovibrio;gut metagenome |
| 0.013717578 | Bacteria;Firmicutes;Clostridia;Clostridiales;Lachnospiraceae |
| 0.013697449 | Bacteria;Firmicutes;Clostridia;Clostridiales;Lachnospiraceae;Oribacterium;uncultured bacterium |
| 0.013144169 | Bacteria;Bacteroidetes;Bacteroidia;Bacteroidales;Prevotellaceae |
| 0.013063153 | Bacteria;Firmicutes;Negativicutes;Selenomonadales;Veillonellaceae;Megasphaera |
| 0.01222598 | Bacteria;Cyanobacteria;Melainabacteria;Gastranaerophilales;uncultured rumen bacterium |
| 0.012052473 | Bacteria;Bacteroidetes;Bacteroidia;Bacteroidales;Prevotellaceae |
| 0.011579542 | Bacteria;Firmicutes;Clostridia;Clostridiales;Lachnospiraceae;Lachnoclostridium |
| 0.011378007 | Bacteria;Bacteroidetes;Bacteroidia;Bacteroidales;Bacteroidaceae;Bacteroides |
| 0.010457905 | Bacteria;Proteobacteria;Deltaproteobacteria;Desulfovibrionales;Desulfovibrionaceae;Desulfovibrio;gut metagenome |
| 0.010244413 | Bacteria;Firmicutes;Clostridia;Clostridiales;Lachnospiraceae;Oribacterium;uncultured bacterium |
| 0.010060483 | Bacteria;Bacteroidetes;Bacteroidia;Bacteroidales;Tannerellaceae;Parabacteroides |
| 0.009949674 | Bacteria;Actinobacteria;Coriobacteriia;Coriobacteriales;Eggerthellaceae;Enterorhabdus;uncultured bacterium |
| 0.009810221 | Bacteria;Firmicutes;Clostridia;Clostridiales;Ruminococcaceae;Oscillibacter |
| 0.009583985 | Bacteria;Actinobacteria;Coriobacteriia;Coriobacteriales;Eggerthellaceae;uncultured |
| 0.009463979 | Bacteria;Firmicutes;Negativicutes;Selenomonadales;Veillonellaceae;Megasphaera |
| 0.009035333 | Bacteria;Firmicutes;Clostridia;Clostridiales;Family XIII |
| 0.008673867 | Bacteria;Firmicutes;Clostridia;Clostridiales;Ruminococcaceae |
| 0.008602365 | Bacteria;Bacteroidetes;Bacteroidia;Bacteroidales;Prevotellaceae |
| 0.008277424 | Bacteria;Firmicutes;Clostridia;Clostridiales;Lachnospiraceae;GCA-900066575;uncultured bacterium |
| 0.008129355 | Bacteria;Firmicutes;Negativicutes;Selenomonadales;Veillonellaceae;Anaerovibrio |

*Continued on next page*

*Table S12 continued*

| Importance | Taxon |
|---|---|
| 0.008123486 | Bacteria;Proteobacteria;Gammaproteobacteria;Betaproteobacteriales;Burkholderiaceae;Sutterella |
| 0.007970951 | Bacteria;Actinobacteria;Coriobacteriia;Coriobacteriales;Eggerthellaceae;Slackia |
| 0.007763456 | Bacteria;Firmicutes;Erysipelotrichia;Erysipelotrichales;Erysipelotrichaceae;Candidatus Stoquefichus |
| 0.00760504 | Bacteria;Verrucomicrobia;Verrucomicrobiae;Verrucomicrobiales;Akkermansiaceae;Akkermansia;uncultured bacterium |
| 0.006748937 | Bacteria;Bacteroidetes;Bacteroidia;Bacteroidales;Rikenellaceae;Rikenellaceae RC9 gut group;uncultured bacterium |
| 0.006731001 | Bacteria;Bacteroidetes;Bacteroidia;Bacteroidales;Prevotellaceae |
| 0.006612061 | Bacteria;Firmicutes;Clostridia;Clostridiales;Lachnospiraceae |
| 0.006583223 | Bacteria;Firmicutes;Clostridia;Clostridiales;Ruminococcaceae;Subdoligranulum |
| 0.006564727 | Bacteria;Firmicutes;Clostridia;Clostridiales;Lachnospiraceae;Oribacterium;uncultured bacterium |
| 0.005657277 | Bacteria;Bacteroidetes;Bacteroidia;Bacteroidales;Prevotellaceae;Alloprevotella;uncultured bacterium |
| 0.005471207 | Bacteria;Firmicutes;Negativicutes;Selenomonadales;Veillonellaceae;Megamonas;uncultured bacterium |
| 0.005405096 | Bacteria;Bacteroidetes;Bacteroidia;Bacteroidales;Bacteroidaceae;Bacteroides |
| 0.005351433 | Bacteria;Bacteroidetes;Bacteroidia;Bacteroidales;Prevotellaceae;Alloprevotella;uncultured bacterium |
| 0.005202657 | Bacteria;Firmicutes;Clostridia;Clostridiales;Lachnospiraceae |
| 0.005201686 | Bacteria;Cyanobacteria;Melainabacteria;Gastranaerophilales |

**Table S13.** Seasonal differences in PICRUSt2-predicted MetaCyc pathway abundances in *Eulemur rufifrons* gut microbiomes (Wilcoxon rank-sum tests). Direction identifies the season with the higher reported pathway abundance.

| Pathway | Dry | Wet | $p$ | $p_{adj}$ | Difference | Direction |
|---|---|---|---|---|---|---|
| PWY-5989 | 0.498628 | 0.878536 | 1.25E-16 | 4.68E-15 | -0.37991 | Higher in Wet |
| PWY0-862 | 0.484984 | 0.837478 | 7.3E-17 | 4.68E-15 | -0.35249 | Higher in Wet |
| PWY-6282 | 0.49277 | 0.843857 | 1.25E-16 | 4.68E-15 | -0.35109 | Higher in Wet |
| PWY-7664 | 0.527624 | 0.872191 | 1.25E-16 | 4.68E-15 | -0.34457 | Higher in Wet |
| PWYG-321 | 0.525207 | 0.814109 | 4.69E-16 | 1.25E-14 | -0.2889 | Higher in Wet |
| FAsynchrony-INITIAL-PWY | 0.416749 | 0.652984 | 4.17E-17 | 4.68E-15 | -0.23624 | Higher in Wet |
| FASYN-ELONG-PWY | 0.70681 | 0.907778 | 3.25E-12 | 3.38E-11 | -0.20097 | Higher in Wet |
| PWY-6147 | 0.577408 | 0.747106 | 8.46E-13 | 1.76E-11 | -0.1697 | Higher in Wet |
| PWY-7539 | 0.600059 | 0.752173 | 1.26E-12 | 2.35E-11 | -0.15211 | Higher in Wet |
| P42-PWY | 0.656167 | 0.807017 | 2.2E-11 | 1.52E-10 | -0.15085 | Higher in Wet |
| PWY-7323 | 0.491628 | 0.641019 | 1.65E-10 | 8.32E-10 | -0.14939 | Higher in Wet |
| PWY-5659 | 0.622144 | 0.768265 | 6.68E-12 | 5.68E-11 | -0.14612 | Higher in Wet |
| PWY-7220 | 0.753421 | 0.898261 | 5.12E-10 | 2.08E-09 | -0.14484 | Higher in Wet |
| PWY-7222 | 0.753421 | 0.898261 | 5.12E-10 | 2.08E-09 | -0.14484 | Higher in Wet |
| NAGLIPASYN-PWY | 0.495013 | 0.63752 | 1.65E-10 | 8.32E-10 | -0.14251 | Higher in Wet |
| PWY-6467 | 0.464664 | 0.603267 | 5.6E-12 | 5.23E-11 | -0.1386 | Higher in Wet |

*Continued on next page*

*Table S13 continued*

| Pathway | Dry | Wet | $p$ | $p_{adj}$ | Difference | Direction |
|---|---|---|---|---|---|---|
| PWY-6700 | 0.620914 | 0.756762 | 2.59E-11 | 1.73E-10 | -0.13585 | Higher in Wet |
| PWY-1269 | 0.486223 | 0.620073 | 2.2E-11 | 1.52E-10 | -0.13385 | Higher in Wet |
| PWY-6703 | 0.524237 | 0.65676 | 2.7E-12 | 2.97E-11 | -0.13252 | Higher in Wet |
| PWY-6737 | 0.835949 | 0.965775 | 5.12E-10 | 2.08E-09 | -0.12983 | Higher in Wet |
| PWY-6612 | 0.561751 | 0.676544 | 4.46E-10 | 1.94E-09 | -0.11479 | Higher in Wet |
| PYRIDNUCSYN-PWY | 0.563658 | 0.677205 | 7.8E-11 | 4.56E-10 | -0.11355 | Higher in Wet |
| PANTOSYN-PWY | 0.605454 | 0.713958 | 4.46E-10 | 1.94E-09 | -0.1085 | Higher in Wet |
| ASPASN-PWY | 0.660713 | 0.768941 | 1.9E-08 | 5.21E-08 | -0.10823 | Higher in Wet |
| POLYISOPRENsynchrony-PWY | 0.588401 | 0.693281 | 2.54E-10 | 1.25E-09 | -0.10488 | Higher in Wet |
| FOLSYN-PWY | 0.615395 | 0.715333 | 8.81E-10 | 3.43E-09 | -0.09994 | Higher in Wet |
| PANTO-PWY | 0.545923 | 0.643163 | 6.65E-09 | 2.15E-08 | -0.09724 | Higher in Wet |
| GLUCONEO-PWY | 0.726125 | 0.821444 | 2.93E-10 | 1.41E-09 | -0.09532 | Higher in Wet |
| PWY-6151 | 0.576513 | 0.664637 | 8.65E-08 | 2.19E-07 | -0.08812 | Higher in Wet |
| PWY-7663 | 0.987005 | 1.073212 | 1.93E-09 | 7.22E-09 | -0.08621 | Higher in Wet |
| PWY-5695 | 0.738091 | 0.822985 | 5.9E-09 | 1.97E-08 | -0.08489 | Higher in Wet |
| PWY-7242 | 0.511883 | 0.588455 | 3.23E-05 | 5.93E-05 | -0.07657 | Higher in Wet |
| CALVIN-PWY | 0.958772 | 1.035025 | 1.07E-08 | 3.22E-08 | -0.07625 | Higher in Wet |
| PWY-6126 | 0.825501 | 0.901479 | 1.07E-08 | 3.22E-08 | -0.07598 | Higher in Wet |
| DTDPRHAMSYN-PWY | 0.678452 | 0.754414 | 4.9E-11 | 2.96E-10 | -0.07596 | Higher in Wet |
| PWY-6897 | 0.580609 | 0.654766 | 1.68E-06 | 3.52E-06 | -0.07416 | Higher in Wet |
| PWY-7229 | 0.854455 | 0.925495 | 2.12E-08 | 5.67E-08 | -0.07104 | Higher in Wet |
| PWY-7199 | 0.470537 | 0.539936 | 7.05E-05 | 0.000123 | -0.0694 | Higher in Wet |
| COA-PWY | 0.772923 | 0.841414 | 7.5E-09 | 2.38E-08 | -0.06849 | Higher in Wet |
| PWY-6123 | 0.842748 | 0.908091 | 2.12E-08 | 5.67E-08 | -0.06534 | Higher in Wet |
| NONMEVIPP-PWY | 0.795087 | 0.859826 | 1.23E-10 | 6.55E-10 | -0.06474 | Higher in Wet |
| PWY-7560 | 0.795087 | 0.859826 | 1.23E-10 | 6.55E-10 | -0.06474 | Higher in Wet |
| PWY-5973 | 0.938587 | 0.999207 | 1.06E-10 | 5.98E-10 | -0.06062 | Higher in Wet |
| COBALSYN-PWY | 0.625833 | 0.686099 | 7E-06 | 1.39E-05 | -0.06027 | Higher in Wet |
| PYRIDNUCSAL-PWY | 0.691813 | 0.750649 | 3.2E-09 | 1.15E-08 | -0.05884 | Higher in Wet |
| 1CMET2-PWY | 0.671162 | 0.728312 | 1.07E-06 | 2.42E-06 | -0.05715 | Higher in Wet |
| SER-GLYSYN-PWY | 0.758735 | 0.812917 | 4.63E-09 | 1.6E-08 | -0.05418 | Higher in Wet |
| PWY-5121 | 0.766512 | 0.817302 | 2.96E-08 | 7.81E-08 | -0.05079 | Higher in Wet |
| THRESYN-PWY | 0.847952 | 0.897819 | 1.2E-08 | 3.56E-08 | -0.04987 | Higher in Wet |
| RIBOSYN2-PWY | 0.700958 | 0.745875 | 3.19E-07 | 7.64E-07 | -0.04492 | Higher in Wet |
| PWY-6609 | 0.708196 | 0.749233 | 0.000262 | 0.000422 | -0.04104 | Higher in Wet |
| PWY-3001 | 0.835671 | 0.87359 | 6.8E-07 | 1.55E-06 | -0.03792 | Higher in Wet |
| PWY-6385 | 0.772555 | 0.810309 | 1.53E-06 | 3.3E-06 | -0.03775 | Higher in Wet |
| TRNA-CHARGING-PWY | 0.777095 | 0.814065 | 2.18E-06 | 4.52E-06 | -0.03697 | Higher in Wet |
| PEPTIDOGLYCANSYN-PWY | 0.778001 | 0.813883 | 1.53E-06 | 3.3E-06 | -0.03588 | Higher in Wet |
| PWY-7221 | 0.757611 | 0.793487 | 1.28E-06 | 2.86E-06 | -0.03588 | Higher in Wet |
| PWY-6387 | 0.787112 | 0.821751 | 3.63E-06 | 7.37E-06 | -0.03464 | Higher in Wet |
| PWY-6386 | 0.783678 | 0.816676 | 3.23E-05 | 5.93E-05 | -0.033 | Higher in Wet |
| PWY-7219 | 0.897355 | 0.929618 | 8.65E-05 | 0.000147 | -0.03226 | Higher in Wet |
| PWY-7208 | 0.920424 | 0.952636 | 0.000179 | 0.000294 | -0.03221 | Higher in Wet |
| PWY-5686 | 0.862465 | 0.892306 | 0.000296 | 0.000474 | -0.02984 | Higher in Wet |
| PWY-2942 | 0.842137 | 0.87159 | 8.08E-05 | 0.00014 | -0.02945 | Higher in Wet |
| PHOSLIPSYN-PWY | 0.768728 | 0.738315 | 5.72E-05 | 0.000102 | 0.030414 | Higher in Dry |
| ARO-PWY | 0.777451 | 0.745183 | 6.14E-05 | 0.000108 | 0.032268 | Higher in Dry |
| PWY-6969 | 0.553271 | 0.518676 | 0.000541 | 0.000851 | 0.034595 | Higher in Dry |
| TCA | 0.462338 | 0.426911 | 0.000453 | 0.000718 | 0.035427 | Higher in Dry |
| COMPLETE-ARO-PWY | 0.813276 | 0.77768 | 1.13E-05 | 2.17E-05 | 0.035596 | Higher in Dry |
| PWY0-162 | 0.678881 | 0.642554 | 4.02E-05 | 7.22E-05 | 0.036327 | Higher in Dry |
| PWY-5101 | 0.903772 | 0.866518 | 7E-06 | 1.39E-05 | 0.037254 | Higher in Dry |

*Continued on next page*

*Table S13 continued*

| **Pathway** | **Dry** | **Wet** | $p$ | $p_{adj}$ | **Difference** | **Direction** |
|---|---|---|---|---|---|---|
| POLYAMSYN-PWY | 0.183304 | 0.143869 | 8.65E-05 | 0.000147 | 0.039435 | Higher in Dry |
| PWY-7184 | 0.392131 | 0.348728 | 2.89E-07 | 7.03E-07 | 0.043403 | Higher in Dry |
| SULFATE-CYS-PWY | 0.158568 | 0.113636 | 0.000574 | 0.000895 | 0.044932 | Higher in Dry |
| OANTIGEN-PWY | 0.488134 | 0.442554 | 2.79E-05 | 5.22E-05 | 0.04558 | Higher in Dry |
| NONOXIPENT-PWY | 1.012027 | 0.966206 | 3.48E-05 | 6.31E-05 | 0.045821 | Higher in Dry |
| PWY0-166 | 0.418474 | 0.369877 | 4.1E-09 | 1.45E-08 | 0.048596 | Higher in Dry |
| PWY-5918 | 0.144917 | 0.09426 | 3.87E-07 | 9.04E-07 | 0.050657 | Higher in Dry |
| GLYCOGENSYNTH-PWY | 0.720094 | 0.667733 | 0.000121 | 0.0002 | 0.052361 | Higher in Dry |
| PWY-6588 | 0.154424 | 0.101259 | 0.000246 | 0.0004 | 0.053166 | Higher in Dry |
| PWY-7187 | 0.423745 | 0.368847 | 3.38E-10 | 1.58E-09 | 0.054898 | Higher in Dry |
| PWY-5154 | 0.329132 | 0.273858 | 1.65E-05 | 3.16E-05 | 0.055274 | Higher in Dry |
| ARG+POLYAMINE-SYN | 0.262591 | 0.206164 | 1.53E-06 | 3.3E-06 | 0.056427 | Higher in Dry |
| PWY-7392 | 0.187506 | 0.129827 | 9.26E-05 | 0.000156 | 0.057679 | Higher in Dry |
| PWY-7197 | 0.418956 | 0.361254 | 3.87E-07 | 9.04E-07 | 0.057702 | Higher in Dry |
| PWY-5188 | 0.323087 | 0.259955 | 1.13E-05 | 2.17E-05 | 0.063132 | Higher in Dry |
| PWY-6478 | 0.159812 | 0.095661 | 7.96E-12 | 6.48E-11 | 0.064151 | Higher in Dry |
| PWY-6353 | 0.162083 | 0.095616 | 2.38E-07 | 5.86E-07 | 0.066467 | Higher in Dry |
| PWY-6901 | 0.168062 | 0.100612 | 3.33E-06 | 6.85E-06 | 0.06745 | Higher in Dry |
| P164-PWY | 0.14843 | 0.080864 | 5.68E-08 | 1.48E-07 | 0.067566 | Higher in Dry |
| PWY0-1261 | 0.400524 | 0.331097 | 0.000113 | 0.000189 | 0.069427 | Higher in Dry |
| SALVADEHYPOX-PWY | 0.157427 | 0.087224 | 1.68E-06 | 3.52E-06 | 0.070203 | Higher in Dry |
| UDPNAGSYN-PWY | 0.403285 | 0.324861 | 1.95E-07 | 4.87E-07 | 0.078424 | Higher in Dry |
| GLYCOLYSIS-E-D | 0.161971 | 0.082092 | 1.51E-08 | 4.34E-08 | 0.079879 | Higher in Dry |
| P23-PWY | 0.171793 | 0.090732 | 1.51E-08 | 4.34E-08 | 0.081061 | Higher in Dry |
| PWY-6608 | 0.183553 | 0.102375 | 6.2E-07 | 1.43E-06 | 0.081178 | Higher in Dry |
| PWY0-1298 | 0.340084 | 0.254221 | 6.32E-08 | 1.62E-07 | 0.085863 | Higher in Dry |
| GALACTUROCAT-PWY | 0.343812 | 0.257315 | 1.9E-08 | 5.21E-08 | 0.086498 | Higher in Dry |
| GALACT-GLUCUROCAT-PWY | 0.279496 | 0.192723 | 7.71E-10 | 3.07E-09 | 0.086773 | Higher in Dry |
| TRPSYN-PWY | 0.64111 | 0.552846 | 3.88E-10 | 1.77E-09 | 0.088264 | Higher in Dry |
| PRPP-PWY | 0.22839 | 0.138044 | 8.89E-06 | 1.75E-05 | 0.090346 | Higher in Dry |
| GLUCUROCAT-PWY | 0.312145 | 0.22151 | 1.69E-08 | 4.8E-08 | 0.090636 | Higher in Dry |
| PWY-5189 | 0.366076 | 0.273415 | 5.9E-09 | 1.97E-08 | 0.09266 | Higher in Dry |
| PWY-5676 | 0.198627 | 0.099983 | 9.5E-09 | 2.96E-08 | 0.098645 | Higher in Dry |
| PWY0-1296 | 0.451778 | 0.348765 | 3.2E-09 | 1.15E-08 | 0.103012 | Higher in Dry |
| PWY-6892 | 0.48661 | 0.378289 | 1.15E-09 | 4.38E-09 | 0.108321 | Higher in Dry |
| P441-PWY | 0.308274 | 0.197532 | 1.98E-16 | 6.18E-15 | 0.110742 | Higher in Dry |
| GLCMANNANAUT-PWY | 0.267271 | 0.155269 | 2.7E-12 | 2.97E-11 | 0.112002 | Higher in Dry |
| PWY0-1297 | 0.326726 | 0.212704 | 3.57E-11 | 2.3E-10 | 0.114022 | Higher in Dry |
| GLUTORN-PWY | 0.519526 | 0.404385 | 4.9E-11 | 2.96E-10 | 0.115141 | Higher in Dry |
| P161-PWY | 0.27549 | 0.159187 | 6.68E-12 | 5.68E-11 | 0.116303 | Higher in Dry |
| DAPLYSINESYN-PWY | 0.570641 | 0.449587 | 1.58E-11 | 1.18E-10 | 0.121054 | Higher in Dry |
| HISTSYN-PWY | 0.590427 | 0.467876 | 5.6E-12 | 5.23E-11 | 0.122551 | Higher in Dry |
| PWY-3781 | 0.251288 | 0.115879 | 2.79E-05 | 5.22E-05 | 0.135408 | Higher in Dry |
| ARGSYNBSUB-PWY | 0.55872 | 0.422474 | 9.47E-12 | 7.38E-11 | 0.136246 | Higher in Dry |
| PWY-7400 | 0.538793 | 0.395543 | 1.53E-12 | 2.38E-11 | 0.143251 | Higher in Dry |
| ARGSYN-PWY | 0.539978 | 0.396616 | 1.53E-12 | 2.38E-11 | 0.143362 | Higher in Dry |
| PWY4FS-7 | 0.57649 | 0.428375 | 1.85E-12 | 2.47E-11 | 0.148114 | Higher in Dry |
| PWY4FS-8 | 0.57649 | 0.428375 | 1.85E-12 | 2.47E-11 | 0.148114 | Higher in Dry |
| PWY-6895 | 0.282636 | 0.119599 | 2.7E-12 | 2.97E-11 | 0.163038 | Higher in Dry |
| PWY-5505 | 0.483035 | 0.296149 | 8.46E-13 | 1.76E-11 | 0.186886 | Higher in Dry |
| PWY-5971 | 0.383279 | 0.176593 | 6.11E-09 | 2.01E-08 | 0.206687 | Higher in Dry |

† $p_{adj}$ = Benjamini–Hochberg false-discovery-rate-adjusted probability. The displayed set has adjusted $p$ <

**Table S11.** Monthly median Firmicutes-to-Bacteroidetes relative abundance ratios in *Eulemur rufifrons* gut microbiomes.

| **Month** | **Median F/B ratio** | **IQR (Q1–Q3)** |
|---|---:|---:|
| January | 0.558 | 0.431–0.732 |
| February | 0.581 | 0.430–0.811 |
| July | 1.210 | 0.893–1.300 |
| August | 0.885 | 0.818–1.190 |
| September | 1.260 | 1.020–1.580 |
| October | 0.475 | 0.350–0.630 |

† F/B ratio = Firmicutes-to-Bacteroidetes relative abundance ratio; IQR = interquartile range.

0.001.

### Supporting Figures

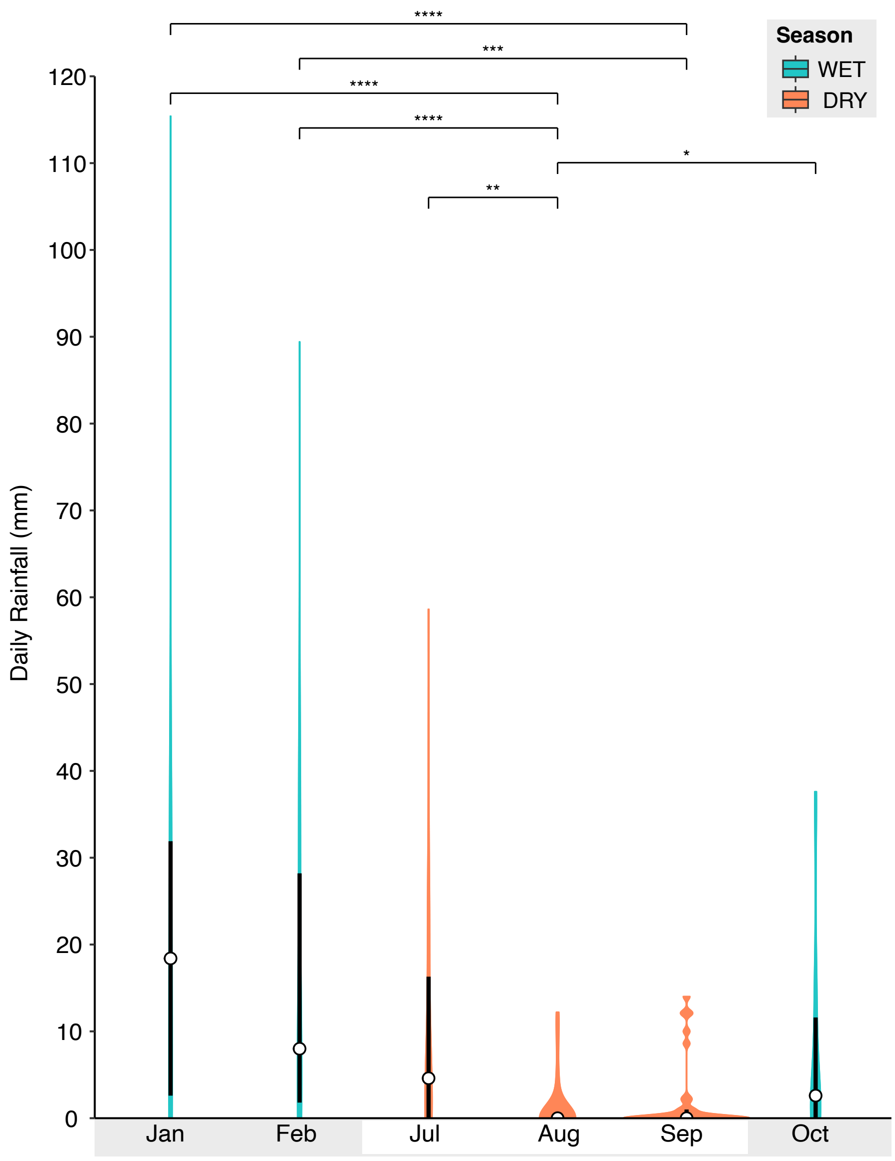


**Figure S1.** Median rainfall across sampling months in Ranomafana National Park in 2020.

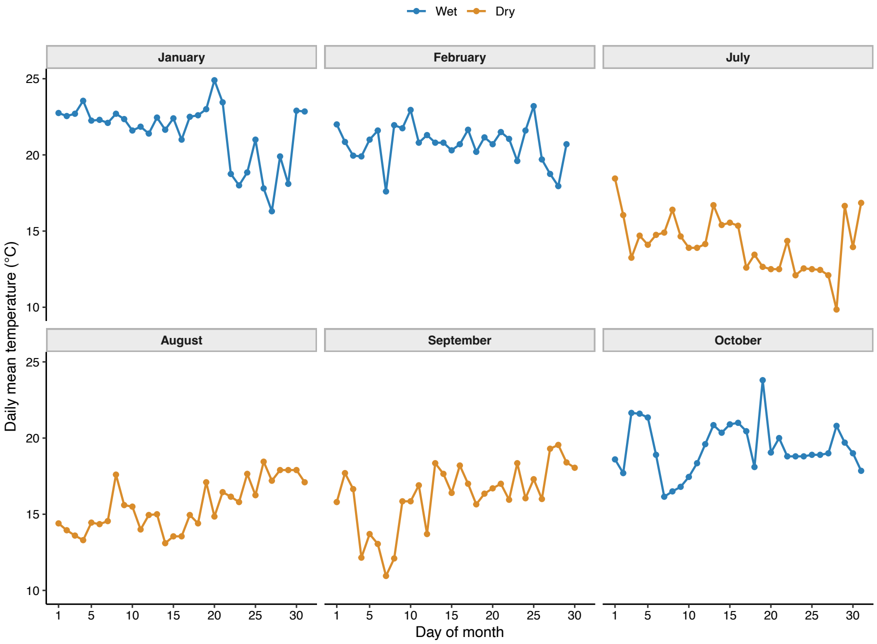


**Figure S2.** Fluctuations in daily temperature across sampling months in Ranomafana National Park in 2020.

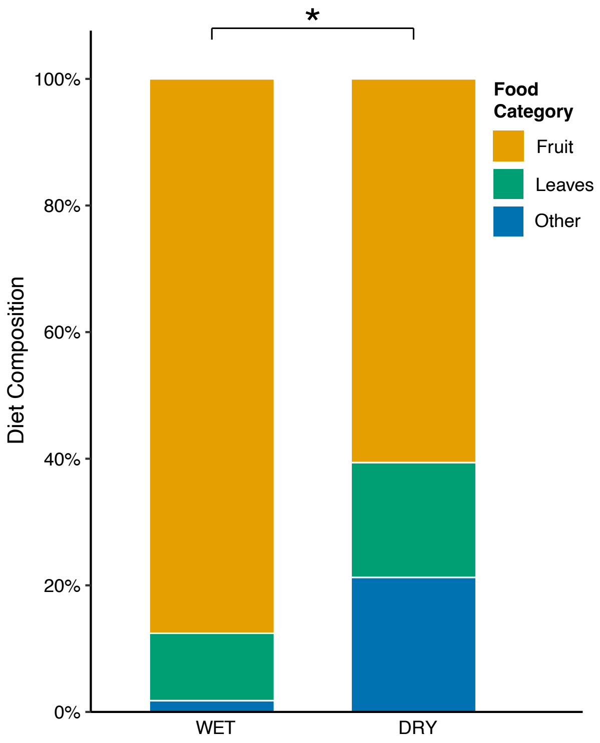


**Figure S3.** Different diet composition of *Eulemur rufifrons* between the wet and dry seasons in 2020, expressed as percentages of feeding observations. Brackets show statistical significance of whole–composition mixed–model tests.

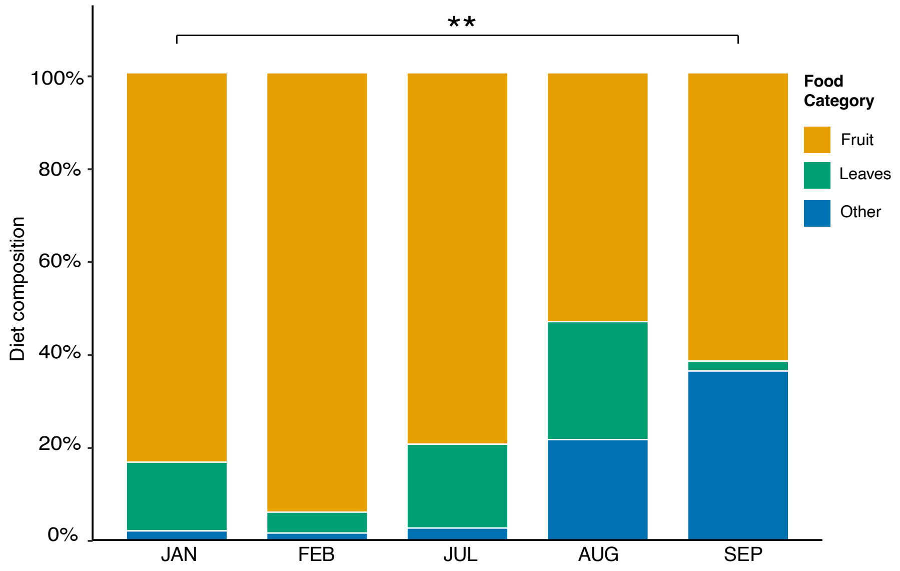


**Figure S4.** Different diet composition of *Eulemur rufifrons* among sampling months in 2020, expressed as percentages of feeding observations. Brackets show overall statistical significance of whole–composition mixed–model tests.

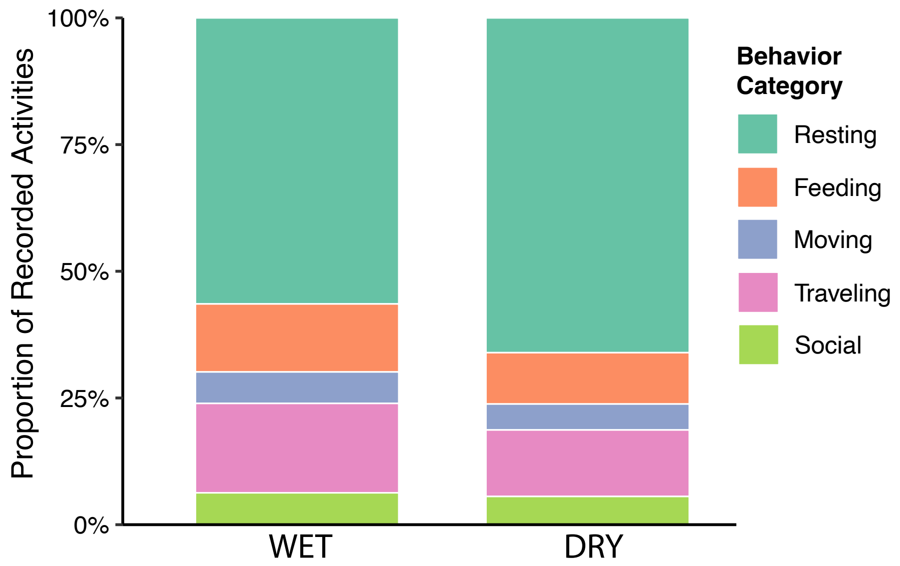


**Figure S5.** Differences in *Eulemur rufifrons* activity budget compositions between the wet and dry seasons in 2020, expressed as percentages of behavioral observations.

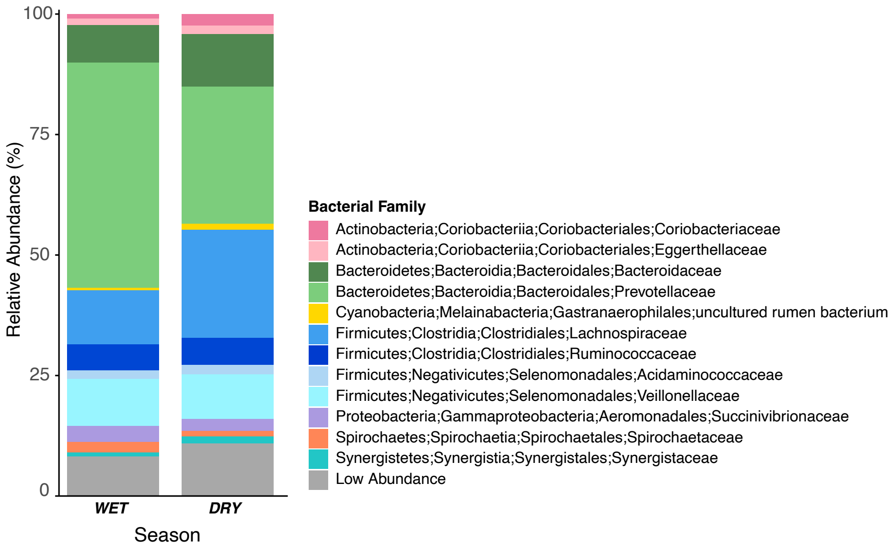


**Figure S6.** Different relative abundances of gut bacterial taxa between the wet and dry season.

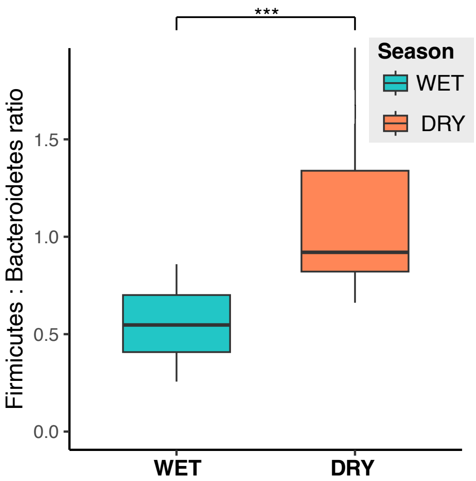


**Figure S7.** Different Firmicutes-to-Bacteroidetes (F/B) relative abundance ratios between the wet and dry season.

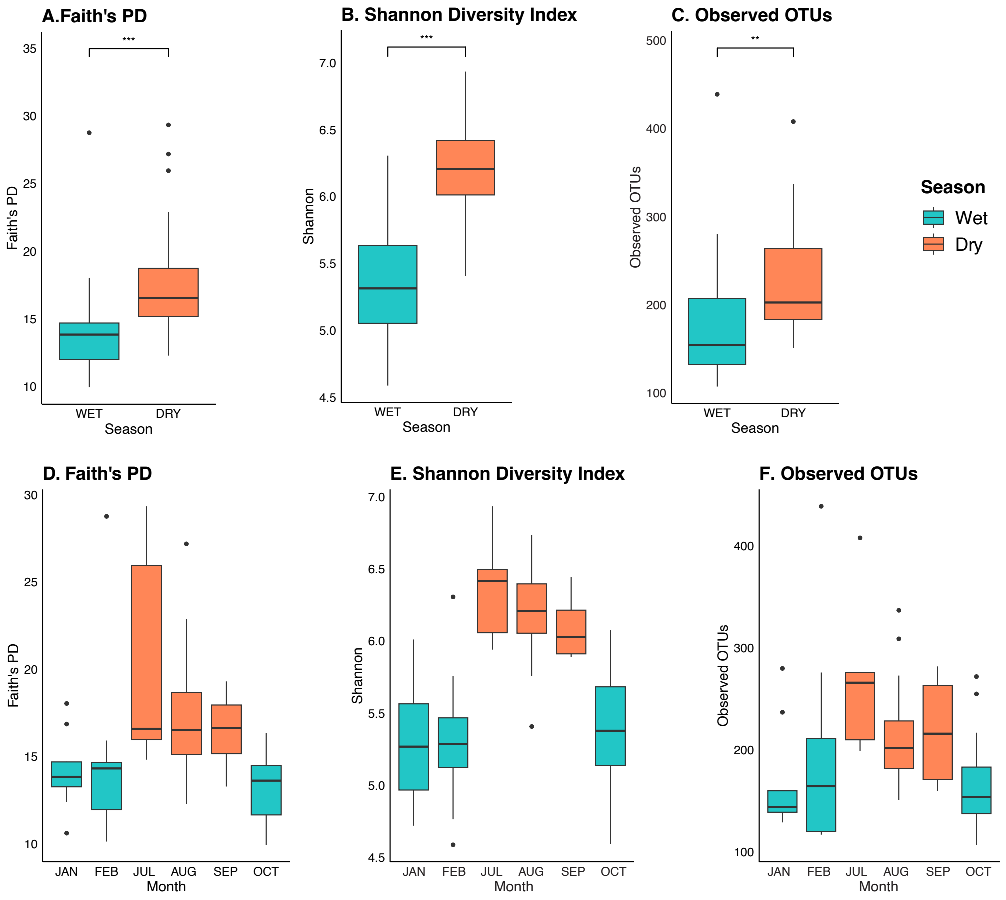


**Figure S8.** Three measures of gut microbiome alpha diversity between seasons and across sampling months: Faith's phylogenetic diversity, Shannon diversity index, and observed operational taxonomic units (OTUs).

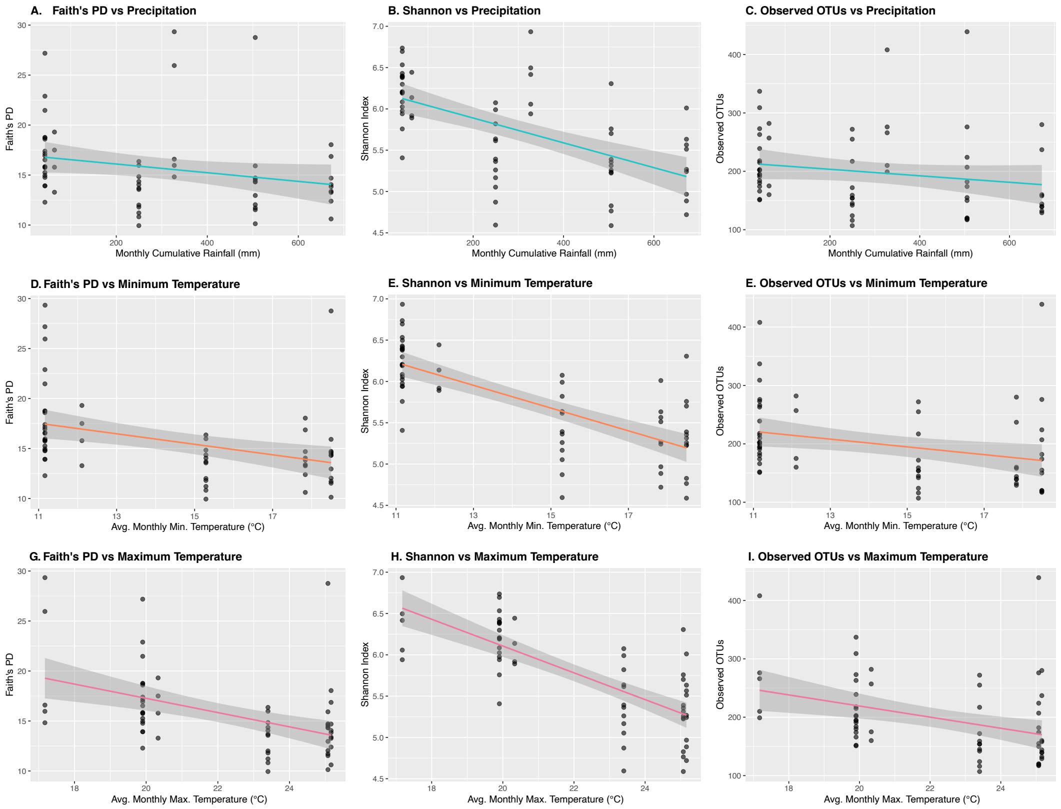


**Figure S9.** Linear associations between three gut microbiome alpha diversity metrics and measured climate variables.

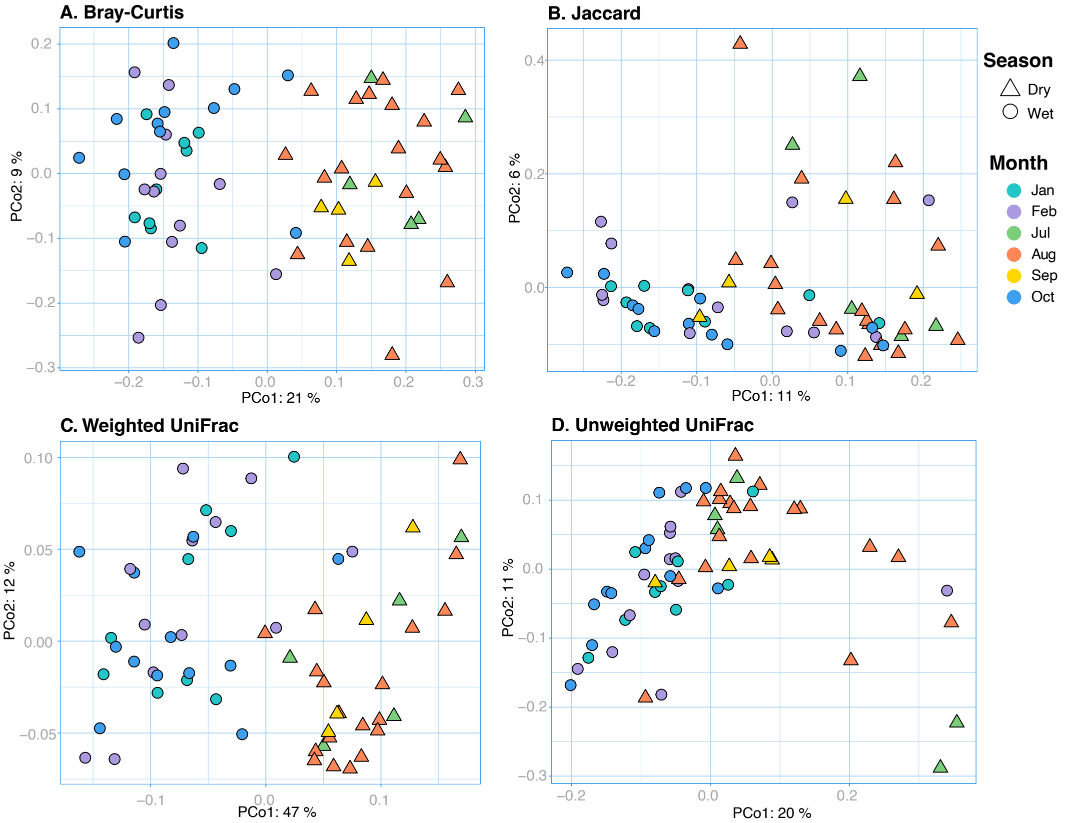


**Figure S10.** Principal coordinates analyses (PCoA) of four gut microbiome beta diversity metrics. More similar samples cluster together in ordination space.

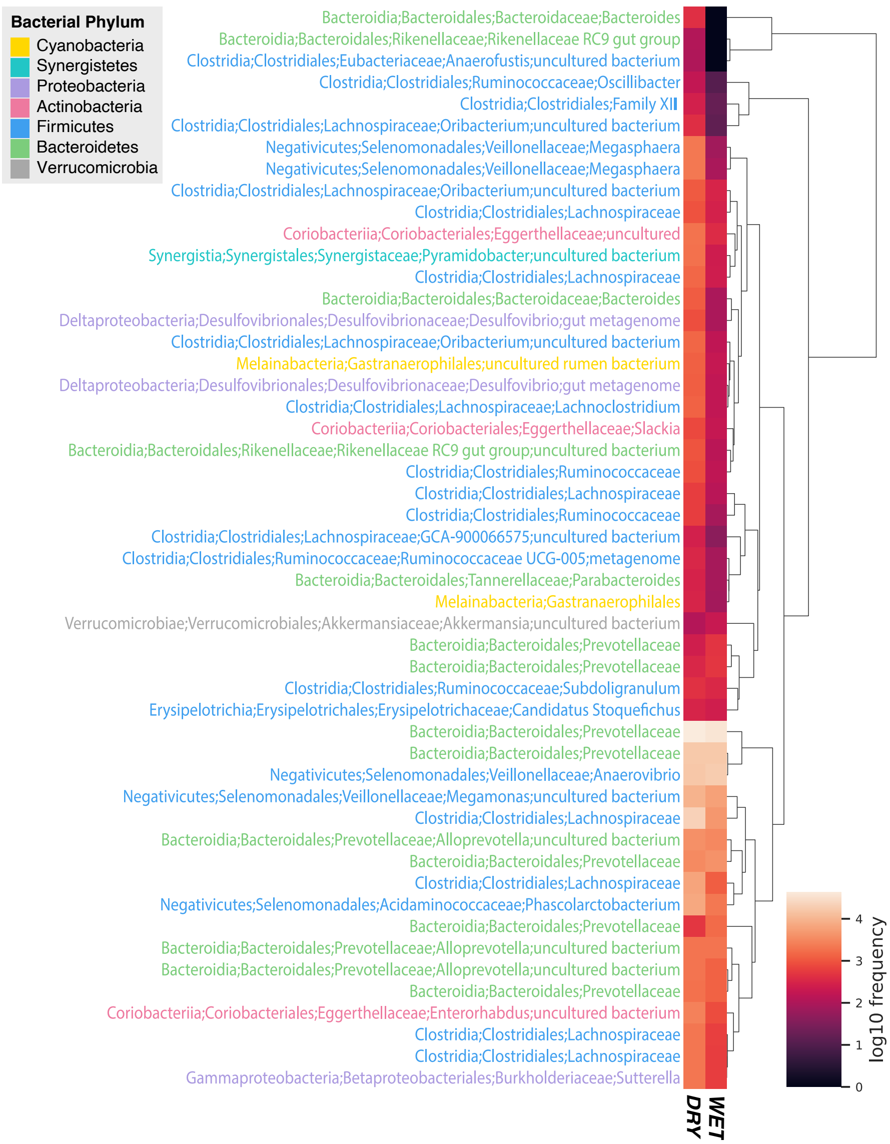


**Figure S11.** Heatmap of the 50 most discriminating microbial taxa in predicting season as a function of microbiome composition. Taxa in Bacteroidetes (green) and Firmicutes (blue) account for 80% of these features. The dendrogram groups taxa by similarity in abundance patterns, with longer branches representing greater dissimilarity.

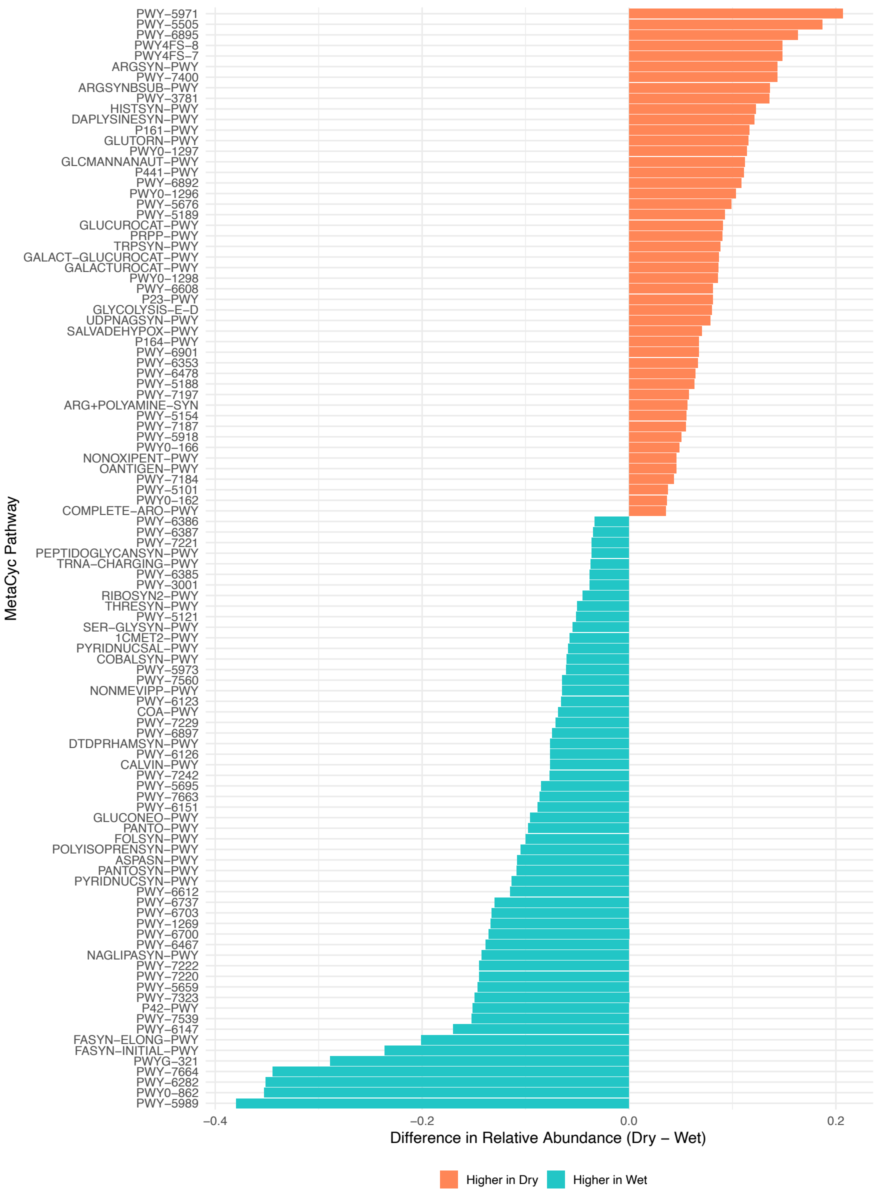


**Figure S12.** Seasonal differences in PICRUSt2-predicted MetaCyc pathway abundances in gut microbiomes of *Eulemur rufifrons*. Bar colors distinguish MetaCyc pathway classes. Displayed pathways have a $p < 0.0001$.
